# DeepPBS: Geometric deep learning for interpretable prediction of protein–DNA binding specificity

**DOI:** 10.1101/2023.12.15.571942

**Authors:** Raktim Mitra, Jinsen Li, Jared M. Sagendorf, Yibei Jiang, Tsu-Pei Chiu, Remo Rohs

**Affiliations:** Department of Quantitative and Computational Biology, University of Southern California; Los Angeles, CA 90089, USA; Departments of Chemistry, Physics and Astronomy, and Computer Science, University of Southern California, Los Angeles, CA 90089, USA

## Abstract

Predicting specificity in protein-DNA interactions is a challenging yet essential task for understanding gene regulation. Here, we present Deep Predictor of Binding Specificity (DeepPBS), a geometric deep-learning model designed to predict binding specificity across protein families based on protein-DNA structures. The DeepPBS architecture allows investigation of different family-specific recognition patterns. DeepPBS can be applied to predicted structures, and can aid in the modeling of protein-DNA complexes. DeepPBS is interpretable and can be used to calculate protein heavy atom-level importance scores, demonstrated as a case-study on p53-DNA interface. When aggregated at the protein residue level, these scores conform well with alanine scanning mutagenesis experimental data. The inference time for DeepPBS is sufficiently fast for analyzing simulation trajectories, as demonstrated on a molecular-dynamics simulation of a *Drosophila* Hox-DNA tertiary complex with its cofactor. DeepPBS and its corresponding data resources offer a foundation for machine-aided protein-DNA interaction studies, guiding experimental choices and complex design, as well as advancing our understanding of molecular interactions.

## Introduction

Transcription factors (TFs) play critical roles in various regulatory functions that are essential to all aspects of life^1^. Therefore, understanding the mechanisms by which proteins target specific DNA sequences is crucial^2^. Extensive research has uncovered myriad binding mechanisms that lead to specific high-affinity binding, including strong electrostatic interaction of arginine in the DNA minor groove^3^, deoxyribose sugar-phenylalanine stacking^4^, bidentate hydrogen bonds between guanine (G) and arginine (Arg) in the major groove,^5^ and other interactions^6–8^.

Protein-DNA structures are typically obtained through X-ray crystallography, nuclear magnetic resonance, or cryogenic electron microscopy experiments and stored in the Protein Data Bank (PDB)^9^. Generally, these structures display one bound DNA sequence and the associated physicochemical interactions^6^, but do not encompass the full range of potentially bound DNA sequences. Conversely, this information can be experimentally obtained through protein binding microarray (PBM)^10^, systematic evolution of ligands by exponential enrichment combined with sequencing (SELEX-seq)^11^, chromatin immunoprecipitation sequencing (ChIP-seq)^12^, or high-throughput SELEX (HT-SELEX)^13^. These experiments capture the range of possible bound DNA sequences but do not necessarily provide structural information. In essence, these sets of experiments are complementary, and manual examination is often required to correlate molecular interaction details from structural data with binding specificity data.

Predicting binding specificity for a given protein sequence, across protein families, remains a challenging and unsolved problem, despite progress for specific protein families^14–21^. Structural changes in the context of binding, along with large mechanistic diversity, contribute to the difficulty^22^. Protein-DNA structures contain valuable information that artificial intelligence can leverage to achieve generalizability across families. In this framework, we introduce Deep Predicter of Binding Specificity (DeepPBS). This deeplearning model is designed to capture the physicochemical and geometric contexts of protein-DNA interactions to predict binding specificity, represented as a position weight matrix (PWM^23^) based on a given protein-DNA structure (Fig. 1a). DeepPBS functions across protein families (Fig. 2) and acts as a bridge between structure-determining and binding specificity-determining experiments.

**Fig. 1.**
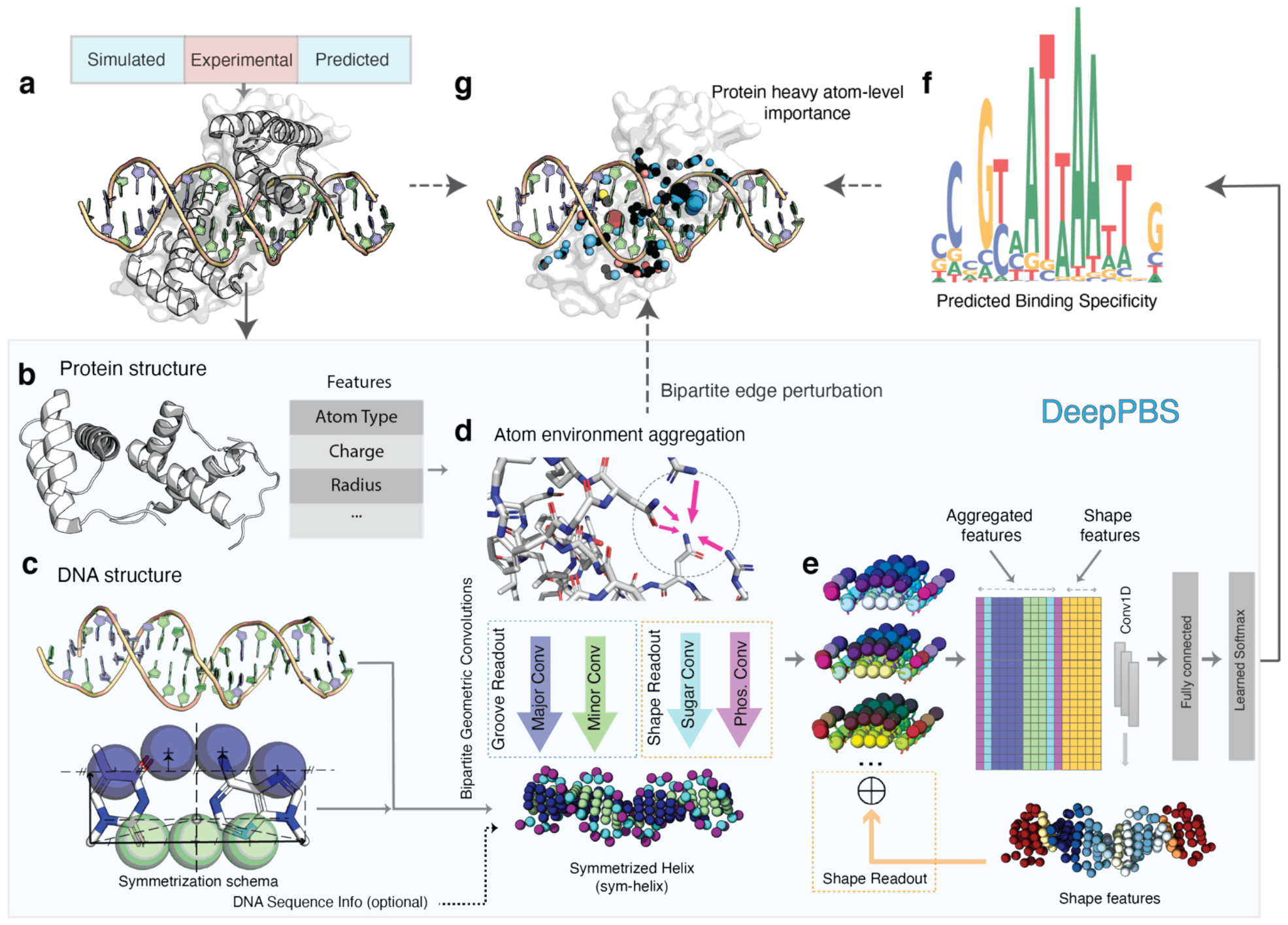
Schematic illustration of the DeepPBS framework. **(a)** DeepPBS input (PDB ID 2R5Y in this example) and possible input sources. **(b)** Protein structure (heavy atom graph, with features computed for each vertex). **(c)** Symmetrization schema in base-pair frame applied to DNA structure, resulting in a symmetrized helix (sym-helix). **(d)** Spatial graph convolution on the protein graph for atom environment aggregation, followed by bipartite geometric convolutions from protein graph vertices to sym-helix points (shown as spheres with specific colors for major groove, minor groove, phosphate, and sugar). **(e)** Three-dimensional sym-helix is flattened with aggregated information (concatenated with computed shape features) into a one-dimensional (1D) representation, followed by 1D convolutions and regression onto base probabilities. **(f)** DeepPBS outputs binding specificity. **(g)** Effect of perturbing bipartite edges involved in **(d)** can be measured in terms of changes in the output, providing an effective measure of interpretability. Abbreviations used in figure: Phos (Phosphate), Conv (Convolutions).

**Fig. 2.**
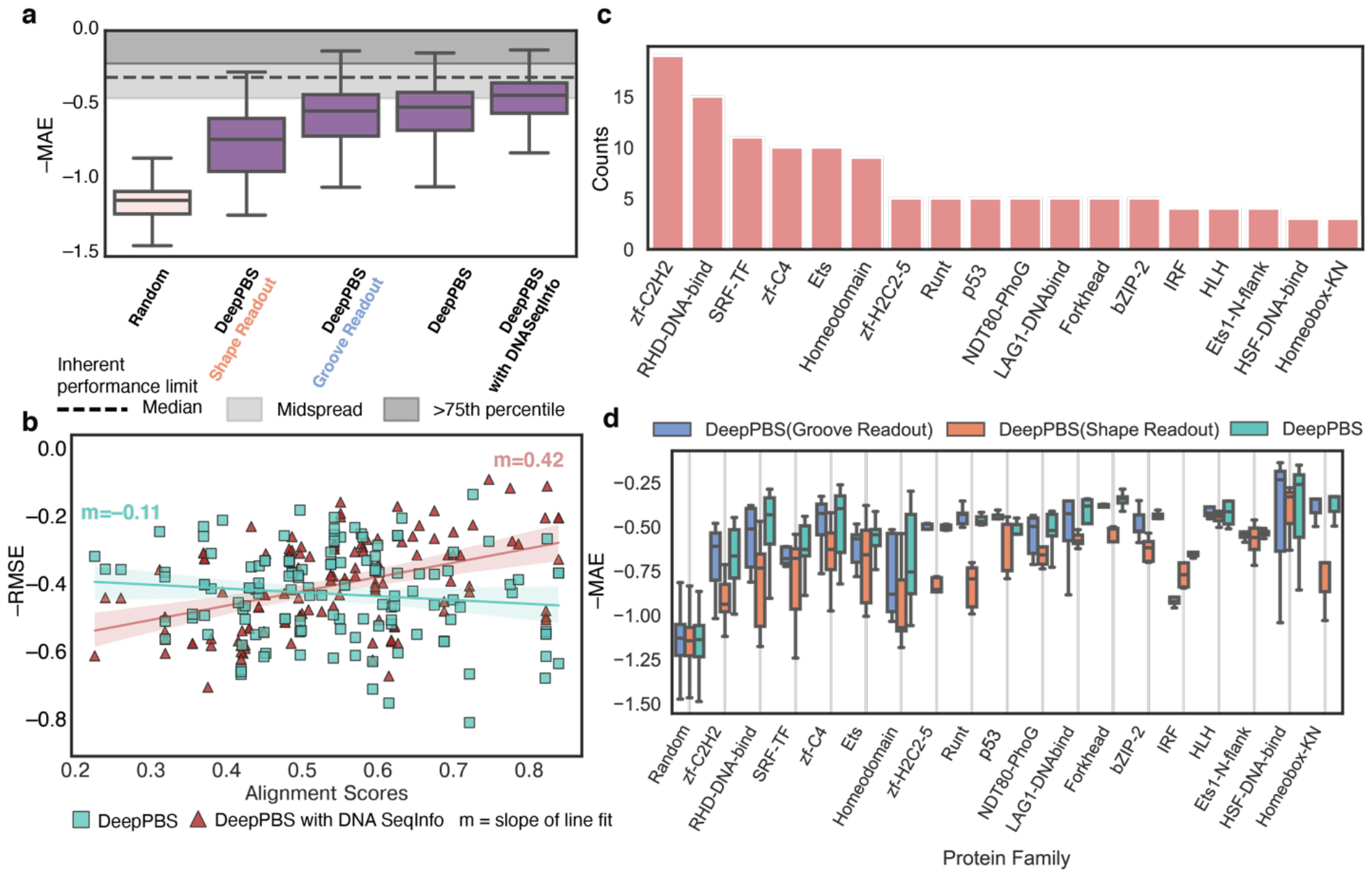
Performance of DeepPBS for predicting binding specificity across protein families on experimentally determined structures. **(a)** Prediction performances of DeepPBS along with “Groove Readout", “Shape Readout” and “with DNASeqInfo” variations, on benchmark set (Supplementary Section 1), with outliers removed. **(b)** Performances of DeepPBS and “with DNASeqInfo” models in context of PWM–co-crystal DNA alignment score (Supplementary Section 2). Shaded regions indicate 95% confidence interval for corresponding linear fit. **(c)** Abundances of various protein families (PFAM annotations) in constructed benchmark set (counts > 3). **(d)** Performances of DeepPBS, “Groove Readout”, and “Shape Readout” models across various protein families (counts > 3), with outliers removed. All benchmark predictions are made by an ensemble average of five models trained via cross-validation. Cross-validation performances of individual trained models are shown in Fig. S5a. For box plots in **(a)** and **(d)**, lower limit represents lower quartile, middle line represents median, and upper limit represents upper quartile. Whiskers do not include outliers.

Input to DeepPBS is not limited to experimental structures (Fig. 1a). The rapid advancement of protein structure prediction models, including AlphaFold^24^ and RoseTTAFold^25^, along with protein-DNA complex modelers, such as RoseTTAFoldNA^26^ (RFNA) and MELD-DNA^27^, have led to an exponential increase in the availability of structural data for analysis. This scenario highlights the growing need for a generalized computational model to analyze protein-DNA structures. We demonstrate how DeepPBS can work in conjunction with structure prediction methods for predicting specificity for proteins without available experimental structures (Fig. 3a-d). In addition, the design of a protein-DNA complex can be improved by optimizing bound DNA using DeepPBS feedback (Fig. 3e-g). We show that this pipeline is competitive with the recent family-specific model rCLAMPS^28^ (Fig. 3h-i) while being more generalizable: specifically, DeepPBS is family-agnostic, can handle biological assemblies, and can predict flanking preferences.

**Fig. 3.**
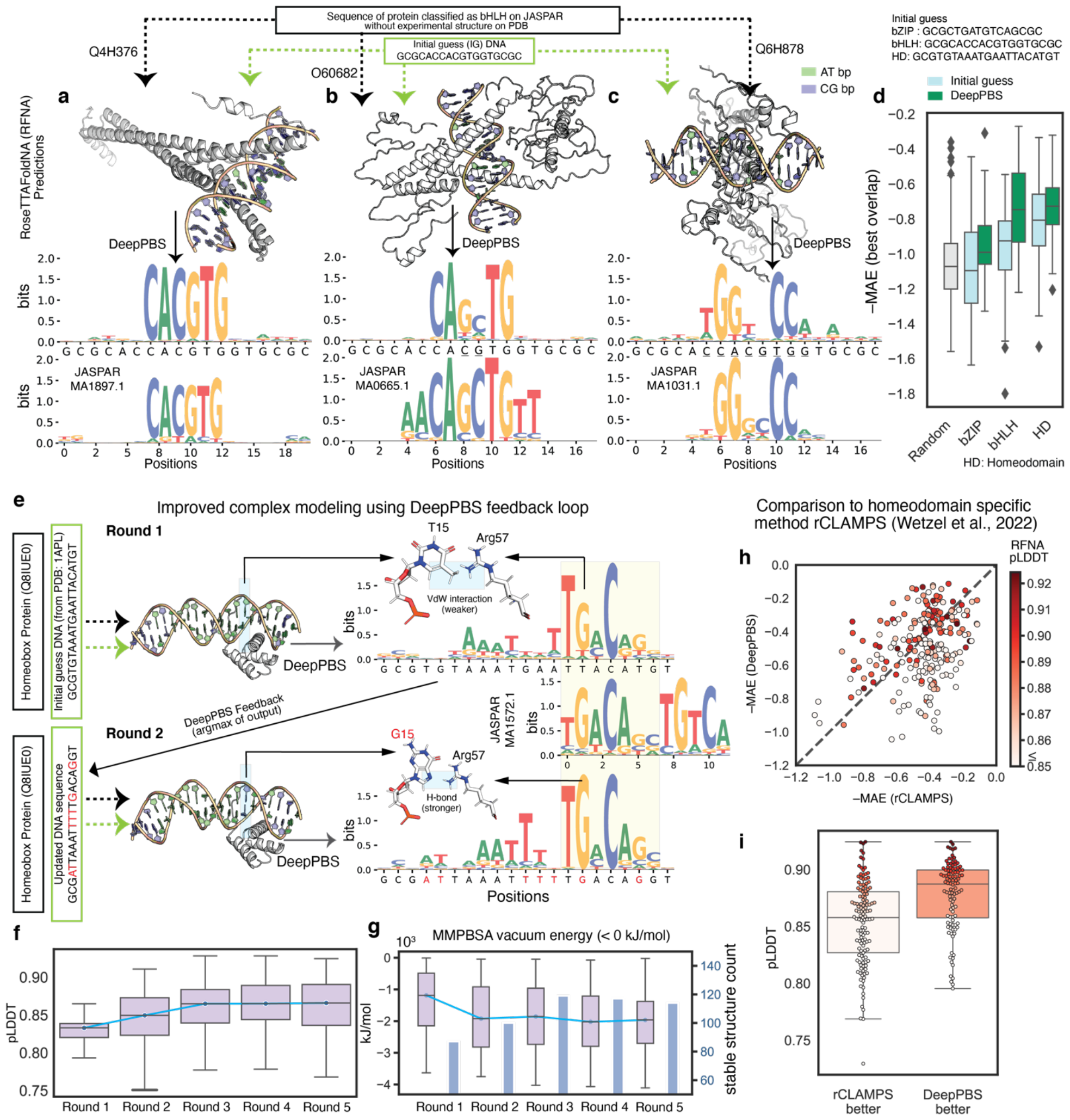
Application of DeepPBS on predicted protein-DNA complex structures. Various predictive approaches (e.g., RFNA, MELD-DNA) can be used to predict protein-DNA complex structures in the absence of experimental data. DeepPBS can predict binding specificity based on this predicted complex. Shown are examples for three full-length bHLH protein sequences: **(a)** Max homodimer from *Ciona intestinalis*, **(b)** MSC dimer from *Homo sapiens*, and **(c)** OJ1581_H09.2 dimer from *Oryza sativa*. **(d)** Performance of DeepPBS via the same process applied for three different families, bZIP (n = 50), bHLH (n = 49), and HD (n = 236), compared to baselines determined for random (drawn from uniform) and IG DNA sequences. Each protein has a unique JASPAR annotation and lacks an experimental structure for the complex. Structures for protein complexes were predicted by RFNA. Proteins passed the pre-processing criterion of DeepPBS. **(e)** One iteration of DeepPBS feedback, demonstrated for human TGIF2LY protein. (vdW: van der Waals). **(f)** RFNA-predicted LDDT^39^ score over rounds 1-5 of DeepPBS feedback loop, with outliers removed. **(g)** Calculated MM-PBSA (Supplementary Section 7) vacuum energy distribution for stable RFNA predictions (< 0 kJ/mol) and corresponding counts over rounds 1-5, with outliers removed. **(h)** Comparison of DeepPBS predictions against HD family-specific method rCLAMPS, color-coded by pLDDT. Diagonal line represents y=x. **(i)** Distribution of pLDDT for two cases: when DeepPBS outperforms rCLAMPS (above diagonal in **(h)**) and vice versa (below diagonal in **(h)**). Box colors denote average pLDDT, using same colormap as in **(h)**. For box plots in **(d), (f), (g)**, and **(i)**, lower limit represents lower quartile, middle line represents median, and upper limit represents upper quartile. Whiskers do not include outliers.

In terms of interpretability, “relative importance” (RI) scores for different heavy atoms in proteins that are involved in interactions with DNA can be extracted from DeepPBS (Fig. 4). As a case study on an important protein for cancer dynamics, we analyze the p53-DNA interface via these RI scores and relate them with existing literature for validation. Additionally, we show that the DeepPBS scores align well with existing knowledge and can be aggregated to produce reasonable agreement with alanine scanning mutagenesis experiments^29^ (Fig. 4i). DeepPBS can be used to analyze simulation trajectories. We demonstrate an example by applying DeepPBS to a molecular dynamics (MD) simulation of an AlphaFold-based modeled complex of the Extradenticle (Exd) and Sex combs reduced (Scr) Hox system^30^ (Fig. 5, Fig. S6, Movie S1).

**Fig. 4.**
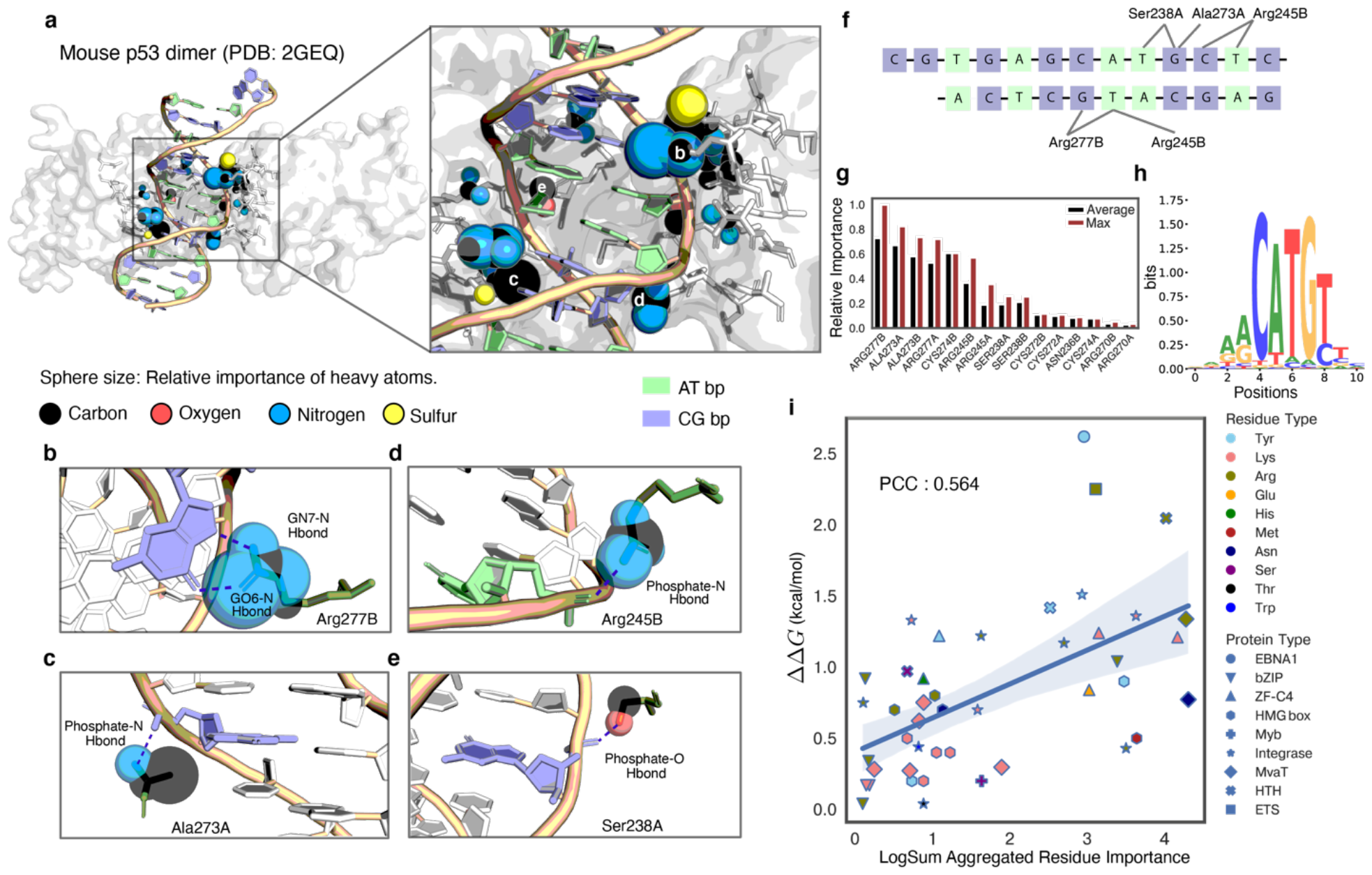
Visualization of DeepPBS importance scores on p53-DNA as a case study, and experimental validation. p53 binds to DNA as a tetramer with two symmetric protein-DNA interfaces^42^. To simplify the biological complex, a p53 dimer structure was chosen^43^ (A and B refer to each monomer, PDB ID: 2GEQ). **(a)** Relative importance (RI) score (i.e., normalized by maximum across atoms) calculated for heavy atoms (denoted by sphere sizes, largest 1, smallest 0) within 5 Å of the sym-helix for a mouse p53 dimer. **(b-e)** Zoomed-in view of specific interactions with different importance scores assigned by DeepPBS. **(f)** Schematic representation of relative arrangement of interactions shown in **(b-e)** along the co-crystal DNA sequence. **(g)** Residue importance computed by average and maximum aggregation of heavy atom importance. **(h)** DeepPBS prediction. **(i)** Comparison of log sum aggregated residue importance computed from DeepPBS ensemble, with experimental free energy change (ΔΔ*G*) determined by alanine scanning mutagenesis experiments. Blue line indicates linear regression fit. Light blue region indicates corresponding 95% confidence interval.

**Fig. 5.**
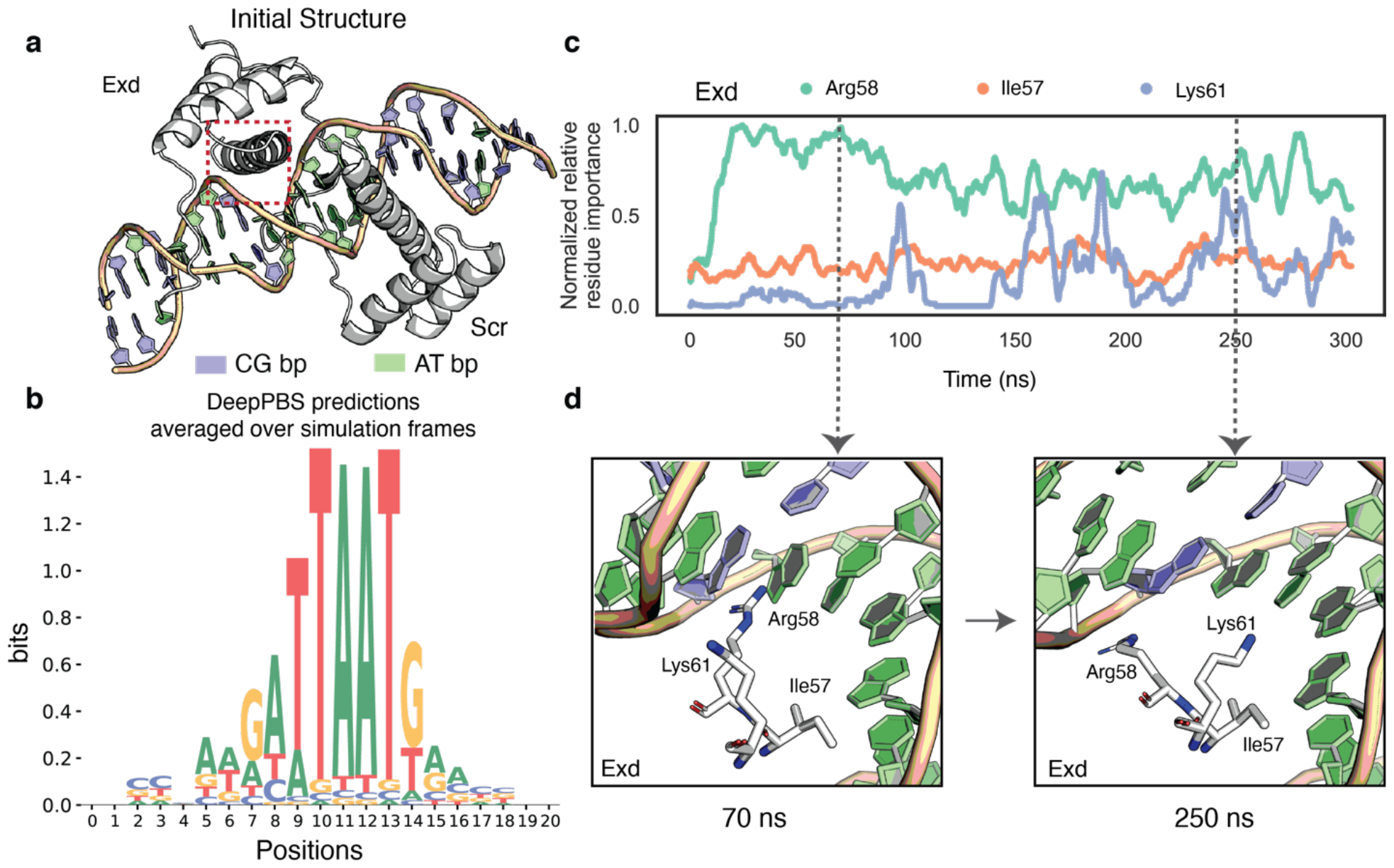
Application of DeepPBS to MD simulation of AlphaFold2- and PDB (2R5Z)-based modeled complex of Exd-Scr-DNA system. (a) Initial structure for the simulation. Red area indicates location of residues of interest (Exd-DNA major groove). **(b)** Average DeepPBS ensemble prediction over 3,000 simulation frames spanning 300 ns. **(c)** Residue-level RI score over time (averaged per 5 ns window) for residues involved in Scr-DNA major groove interaction. **(d)** Snapshot of interactions by Exd Arg58, Lys6, and Ile57 at 70 ns and 250 ns.

### The DeepPBS Framework

The DeepPBS framework is illustrated in Fig. 1. Input to DeepPBS (Fig. 1a) is comprised of one protein-DNA complex structure, with one or more protein chains bound to a DNA double helix. Potential sources for such structures include experimental data (e.g., PDB^9^), simulation snapshots, or designed complexes. DeepPBS processes the structure as a bipartite graph with distinct spatial graph representations for protein and DNA components. The protein graph is an atom-based graph, with heavy atoms as vertices. Several features are computed on these vertices (Fig. 1b). Further information on protein representation and feature computation is available in Supplementary Section 4. We represent DNA as a symmetrized-helix (sym-helix), as detailed in the Methods. This representation removes any sequence identity that the DNA possesses, while preserving the shape of the DNA helix, a critical aspect for protein-DNA binding^3^. Optionally, DNA sequence information can be reintroduced as a feature on the sym-helix points.

DeepPBS performs a series of spatial graph convolutions on the protein graph to aggregate atomic neighborhood information (Fig. 1d). The next, crucial component of DeepPBS consists of a set of bipartite geometric convolutions applied from the protein graph to the sym-helix (Fig. 1d). Strong chemical interactions (e.g., hydrogen bonds) depend on both location and orientation^5^. DeepPBS learns how the geometric orientation of the sym-helix points is associated with the orientations and chemistry of neighboring protein residues. Four distinct bipartite convolutions are employed for the sym-helix points, corresponding to the major groove, minor groove, and the phosphate and sugar moieties. Major and minor groove convolutions are referred to as “Groove Readout”. Phosphate and sugar moiety convolutions, combined with DNA shape information, form the “Shape Readout” (Fig. 1e). The “Groove Readout” and “Shape Readout” factors collaboratively determine binding specificity to varying extents for different protein families. At this point, the sym-helix representation enables a straightforward flattening of aggregated features on the three-dimensional (3D) sym-helix to the one-dimensional (1D) base pair-level features. By adding DNA shape information and implementing 1D convolutional neural network (CNN) and prediction layers (Fig. 1e), DeepPBS ultimately predicts binding specificity (Fig. 1f). Further architectural details are described in Supplementary Section 5.

Lack of an existing published standard dataset for predicting binding specificity across families from protein-DNA complex structure data made it necessary for us to build a dataset for cross-validation and benchmarking. Details of this process can be found in the Methods section.

### Performance of DeepPBS on experimentally determined structures from PDB

The DeepPBS ensemble (Methods) was employed to evaluate model performance against a benchmark set, as outlined in Supplementary Section 1. The DeepPBS architecture allows models to be trained on two mechanisms: “Groove Readout”, which does not involve backbone convolutions and excludes shape information, and “Shape Readout”, which does not involve groove convolutions (Fig. 1d-e). Benchmark performances of “DeepPBS” (performs both readout modes) and these two variations are shown in Fig. 2a. The performance of the “DeepPBS” model surpassed that of either component in isolation (Fig. 2a). The dataset was constructed using experimentally determined structures; thus, the co-crystal DNA sequence typically serves as a reasonable example of a bound sequence. As expected, integrating sequence information into the sym-helix points (“DeepPBS with DNA SeqInfo”) enhanced performance (Fig. 2a), significantly closing the gap towards the inherent performance limit in the dataset (Supplementary Section 1). However, from both the interpretability and design perspectives, particularly when the bound DNA sequence may not be representative, the “DeepPBS” model is optimal due to its low sensitivity to the DNA sequence in the structure. This fact is evidenced by comparing performances of the “DeepPBS” and “DeepPBS with DNA SeqInfo” models in the context of the PWM–co-crystal DNA alignment score (Supplementary Section 1). Compared to the line fitted to the variation with DNA sequence information (slope = 0.42), the slope of the line fitted to the DeepPBS predictions was closer to zero (Fig. 2b).

As an example, we show the DeepPBS ensemble prediction on the human nuclear factor kappa-light-chain-enhancer of activated B cells (NF-κB) biological assembly from the benchmark dataset. Although the co-crystal DNA sequence was not of the highest affinity, as indicated by experimental data from HOCOMOCO^31^, our prediction circumvented this issue, predicting a binding specificity that was more closely aligned with the experimental data (Fig. S5d). Similar trends (Fig. S5a-c) can be observed from cross-validation predictions by individual DeepPBS models (see Methods). We also included example DeepPBS ensemble predictions (Fig. S7) for structures in the PDB that correspond to specific interactions but do not have a PWM in the two binding specificity databases considered (see Methods). In addition, example DeepPBS ensemble predictions (Fig. S8) for structures of nonspecific protein-DNA binding (e.g., SSO7D-DNA interaction^32^) present in the PDB are presented. These predictions have notably lower information content (IC) compared to those in Fig. S7.

### DeepPBS captures information about various family-specific binding modes

Abundances of different protein families in the benchmark set are described in Fig. 2c (Fig. S5b for crossvalidation set). Family annotations were obtained from the Database of Protein Families (PFAM)^33^. The dataset encompasses a wide range of DNA binding protein families. Performance of DeepPBS on various protein families provides several key insights. DeepPBS showed reasonable generalizability across families, performing well even for families with relatively fewer structures (Fig. 2d, Fig. S5c), such as heat shock factor (HSF) proteins. This observation suggests that the model is learning the underlying mechanisms of protein-DNA binding rather than overfitting on family-specific patterns.

Further validation is provided by comparing performances of the DeepPBS “Groove Readout” and “Shape Readout” models (Fig. 2d, Fig. S5c). For families like zf-C2H2, zf-C4, and HLH, the “Shape Readout” model did not perform as well as the “Groove Readout” model, whereas the full model (“DeepPBS”) performed as well as or slightly better than the “Groove Readout” model. This result aligns with the common understanding of the binding mechanism of these families. For example, zf-C2H2 uses a recognition *α*-helix to scan DNA for suitable base interactions, with minimal DNA bending or conformational change^34^. This binding mode makes the zf-C2H2 family a popular target of protein sequence-based binding specificity prediction and design^14,16,17,21,35^. Conversely, families like IRF (Fig. 2d, Fig. S5c) and T-box (Fig. S5c) showed higher performances for the “Shape Readout” model, consistent with their known binding mechanisms that involve significant conformational changes^4,36^. For families such as Ets (Fig. 2d, Fig. S5c), the “DeepPBS” model outperformed both the “Groove Readout” and “Shape Readout” components. This result suggests that the network captures complex higher-order relationships of these components.

### Application of DeepPBS on computationally predicted protein-DNA complexes

The DeepPBS framework is not limited to experimental structures. Recent advances in scalable structural prediction approaches, driven by artificial intelligence^24,25^, offer unprecedented potential. Specifically, models like RFNA^26^ and MELD-DNA^27^ can be used to predict the structures of protein-DNA complexes from sequences. Such prediction algorithms have paved the way for DeepPBS to be applicable to proteins that lack experimental DNA-bound structure data.

We suggest one potential approach for working with predictive structures in DeepPBS. First, we come up with an initial guess of the DNA (IG DNA) sequence bound to each protein of interest based on the protein family consensus. Then, we use RFNA to predict the protein-DNA bound complex structure, followed by DeepPBS to predict binding specificity. We demonstrate this process (Fig. 3a-c) for three proteins classified as bHLH (basic Helix Loop Helix) in JASPAR^37^. In all three cases, the PDB lacked experimental protein-DNA complex structure. The IG DNA (see Supplementary Section 7) has an E-box motif (“CACGTG”) in the center, which is known^38^ to be a bHLH family target. The first example (UniProt Q4H376, Fig. 3a) is a Max homodimer, for which DeepPBS predicted a specificity closely mirroring that of the IG DNA. The second example (MSC dimer, O60682) was more complicated; the central “CACGTG” in the IG DNA was erroneously assumed, yet DeepPBS successfully predicted the correct motif as “CAGCTG” (Fig. 3b). The third example (Fig. 3c, protein OJ1581_H09.2, Q6H878) does not conform to any E-box motif. Nevertheless, DeepPBS predicted a binding specificity closely mirroring the experimental data (Fig. 3c).

We ran the DeepPBS pipeline for full-length UniProt protein sequences, each with a unique JASPAR entry and no experimental structure for the complex, across three different families (see Supplementary Section 7): bZIP, bHLH, and homeodomain (HD) families. DeepPBS predictions based on RFNA predicted structures exhibited an improved –MAE (i.e., closer to experimental data) compared to the IG DNA baseline (Fig. 3d). An application of DeepPBS to a MELD-DNA predicted complex of mouse CREB1 protein is demonstrated in Fig. S9b. Thus, DeepPBS can take predicted structures from suboptimal DNA sequences and predict specificity close to experimental data.

We next explored whether DeepPBS prediction could be used as feedback (in a loop) to enhance modeling of the protein complex (and, subsequently, improve DeepPBS prediction). We demonstrated this process for the human TGIF2LY protein (UniProt ID Q8IUE0, unstructured region trimmed, Supplementary Section 7) in Fig. 3e. In round 1, we applied RFNA to this protein sequence alongside the IG DNA sequence for the HD family, and then inputted the predicted complexes into DeepPBS. For IG DNA position T15 (Fig. 3e, round 1), DeepPBS predicted a strong preference for G. In the round 1 RFNA output, Arg57 and T15 were involved in one H-bond and one van der Waals interaction. These interactions are theoretically inferior to (weaker than) the possible double H-bonds between a G and Arg57. In round 2, we altered the RFNA input by taking the argmax (the most preferred sequence) from the DeepPBS output (Fig. 3e, round 2). The subsequently folded structure reflected a more robust double H-bond interaction between G15 and Arg57, with the DeepPBS prediction more closely aligning with the experimental data (note positions (round 2) A18, G19, and G20, corresponding to positions 4-6 in MA1572.1; Fig. 3e).

We repeated this DeepPBS prediction process for a total of five rounds, for the set of HD monomer sequences (Supplementary Section 7). The RFNA-predicted confidence metric (pLDDT, LDDT^39^ reflects similarity between the predicted and reference structure for a complex; Supplementary Section 7) improved over these rounds (Fig. 3f). To independently evaluate structure quality, we calculated the Molecular Mechanics and Poisson–Boltzmann Surface Area (MM-PBSA^40^) binding energy (Supplementary Section 7). From round 1 to round 3+, the number of stable structures (binding energy < 0 kJ/mol) increased (Fig. 3g), while their binding energy distributions shifted towards lower values (Fig. 3g). DeepPBS performance improved across the five rounds (Fig. S9a).

The DeepPBS approach to predicting binding specificity fundamentally differs from that of existing methods, which predict specificity based solely on protein sequence information. As a result, comparisons with existing family-specific methods that operate exclusively on protein sequence are unfeasible. However, in conjunction with a complex structure prediction method, we can start from protein sequence information alone and predict binding specificity using DeepPBS. This process can be compared to the recent HD family-specific method, rCLAMPS^28^ (see Supplementary Section 7). rCLAMPS can predict core 6mer specificities for monomer HD proteins. A comprehensive view of performances is presented in Fig. 3h. For significant portions of the data points, DeepPBS outperformed rCLAMPS, and vice versa. DeepPBS generally performed better for higher pLDDT scores, whereas rCLAMPS did better for lower pLDDT scores (Fig. 3i). Thus, the DeepPBS pipeline is comparable to rCLAMPS, while having wider applicability across families, biological assemblies, as well as not being limited to predicting the core binding region alone.

### DeepPBS assessment of protein residue importance at p53-DNA interface

The DeepPBS architecture permits intentional activation or deactivation of specific edges in the bipartite geometric convolution stage (Fig. 1d, Fig. S4). Perturbing a set of edges in this manner will alter the network-predicted result. The mean absolute difference between the original and altered prediction can be used (with proper normalization) as a quantification of the impact of the perturbed set of edges in determining specificity (Fig. 1g, Fig. S4, Methods).

We present results for perturbing edge sets for individual protein heavy atoms (which can also be aggregated to compute residue-level importance). As an example, we examined the protein-DNA interface of p53, a protein crucial for regulating cancer and cell apoptosis^41^. p53 binds to DNA as a tetramer with two symmetric protein-DNA interfaces^42^. To simplify the biological complex, we chose a p53 dimer structure^43^ (PDB ID: 2GEQ). We show the RI scores (with min-max normalization applied) calculated for heavy atoms within 5 Å of the sym-helix (Fig. 4a). Sphere sizes in Fig. 4a denote computed RI scores, with the largest being 1 and smallest 0. The network deems Arg277^33^-guanine bidentate H-bonds (Arg280 in human^44^) as the strongest driver of specificity^5^ (Fig. 4b). Ala273 (Ala276 in human, known for causing apoptosis upon mutation^45^) appears as the second most important driver. Ala273 is involved in H-bonding with a phosphate moiety, using its backbone nitrogen and van der Waals contacts with the DNA major groove (Fig. 4c). This residue has been shown to be an important driver of specificity via van der Waals contacts with the methyl group of thymine in the major groove^44^.

Another important residue according to DeepPBS, Arg245 (Arg248 in human^46^), is present at the DNA minor groove and forms H-bonds with the backbone (Fig. 4d). This decision by the model is primarily based on the orientation of arginine relative to the sym-helix, which is devoid of DNA sequence information. Arg248 is attracted through enhanced negative electrostatic potential due to a narrowing of the minor groove where it binds^42^. Fig. 4e shows the importance assigned to further backbone contacts by Ser238 residues (Ser241 in human). This residue is known^46^ to be important for stabilizing Arg245. The binding specificity prediction of DeepPBS (Fig. 4h) aligns well with known binding patterns of p53, which follows the form RRRC(A/T)(A/T)GYYY (R denotes purine, Y denotes pyrimidine). The interactions shown here are deemed^41,47^ as significant drivers of p53 binding.

### Comparison of DeepPBS-derived residue-level importance with experimental data

We next asked whether DeepPBS-derived importance scores, which reflect the degree to which an interaction determines output binding specificity, can be considered as reliable and potentially physically significant. Although high-affinity interactions can be nonspecific^32,48^, interactions that contribute to high specificity would be expected to maximize affinity across different DNA base possibilities. Therefore, the DeepPBS importance scores associated with these interactions should display some correlation with the corresponding binding affinities. We can test this hypothesis experimentally by using alanine scanning mutagenesis data (Supplementary Section 1). Sets of such experimental data have been made available through recent contributions^49^ in the field. Utilizing these data ^43^, we applied suitable filtering for our context and calculated the log sum aggregated residue level importance scores using DeepPBS (Methods).

A regression plot and Pearson’s correlation coefficient (PCC), as shown in Fig. 4i, illustrate the correspondence between computed values and experimental ΔΔ*G* values for a diverse array of proteins and residues within the protein-DNA interface (Table S1). The obtained PCC of 0.564 corroborates our hypothesis. These results highlight the potential of the DeepPBS computational model as an economical guide for experimentalists who are selecting mutagenesis experiments to conduct at the protein-DNA interface.

### Application of DeepPBS over MD simulation trajectory of Exd-Scr-DNA complex

Owing to a fast inference time, DeepPBS can be used to analyze simulation trajectories. We demonstrated how the protein heavy atom-level interpretability allows automatic detection of conformational changes in the protein-DNA interface. We applied DeepPBS to an MD simulation of the well-studied Exd-Scr-DNA system (Fig. 5a) (PDB ID: 2R5Z)^50–53^. Details of the simulation method are provided in Supplementary Section 8. By computing the DeepPBS prediction over the trajectory, we found that the calculated average ensemble prediction (Fig. 5b) was consistent with the known binding specificity^50^ of the system. Aggregated RI scores for key protein residues interacting in the Exd-DNA major groove are shown in Fig. 5c.

Initially, Arg58 is involved in a bidentate H-bond with G7, which results in a very high RI (Fig. 5c-d, left), whereas Lys61 does not strongly interact. Later, Arg58 rotates away, forming only one H-bond with G7, which allows Lys61 to come closer and interact intermittently (Fig. 5c-d, right), reflecting corresponding changes in RI. Ile57 is involved in a weaker van der Waals interaction throughout the trajectory, resulting in a lower but consistent RI (Fig. 5c-d). Supplementary Section 9 further describes the DeepPBS RI values for residues involved in the Exd-DNA flank and Scr-DNA minor groove interactions. DeepPBS ensemble prediction and heavy atom-level RI scores over the trajectory can be visualized as a movie (Movie S1), which shows how conformational variation of the complex influences the network output. DeepPBS can be used for fast automatic detection of these conformational changes and their effect towards binding specificity determination.

## Discussion

Computationally identifying which DNA sequence, a given protein will bind to remains a challenging question. Although proteins from certain DNA binding families, such as HDs^15,20,54–56^ and C_2_H_2_ ZFs^14–16,18,34,57^, have been studied extensively in this regard, a generalized model of binding specificity remains elusive. This complexity emanates, in part, from the pivotal role that the protein and DNA conformation or shape play in the context of binding specificity. For example, TBX5 undergoes an *α*-helix to 3_!”_-helix conformational change when interacting with DNA. Despite the energy penalty, this transformation, in conjunction with an appropriately matching DNA shape, instigates a strong phenylalanine-sugar ring stacking, thereby facilitating binding^4^. Another example is the Trp repressor protein, which exhibits an almost entirely geometry-driven binding specificity. This protein only makes direct and water-mediated H-bonds with the backbone phosphates^58^, and the DNA shape required for optimal binding gives rise to sequence specificity. Capturing such interactions and how they lead to binding specificity with protein information alone is complicated and cannot be understood in a sequence space alone^22,59^. Furthermore, for many protein families, the protein monomer is insufficient^60^ for binding; a biological assembly, potentially with other interaction partners^61^, is often necessary.

DeepPBS achieves generality across families with the tradeoff of requiring a docked sym-helix, representing a significant step towards solving the larger unsolved problem. As demonstrated in this work, coupling DeepPBS with attempts to model protein-DNA complexes provides a significant step forward in predicting binding specificity across families, based solely on protein information.

DeepPBS allows exploration of exciting future possibilities, including the creation of DNA-targeted protein designs that could potentially contribute to therapeutic advancements. DeepPBS could serve as a preliminary screening tool for devised candidate complexes, ensuring their specificity to the intended target DNA sequence prior to any costly experimental validations. Moreover, recent studies have shown that TF-DNA binding can energetically favor mismatched base-pairs^62^. Given the combinatorial complexity of possible hypotheses, deciding which mismatch experiments to perform to discover more such instances poses a significant challenge. Although there is currently a lack of training data for mismatches, the DeepPBS architecture, in theory, could facilitate the prediction of mismatched binding specificity. This approach could assist in deciding which experiments to conduct.

In summary, we have introduced a computational framework that distills the intricate structural nuances of protein-DNA binding and bridges this understanding with binding specificity data, effectively connecting structure-determining and specificity-determining experiments. The DeepPBS architecture allows inspection of family-specific “Groove Readout” and “Shape Readout” patterns and their effects on binding specificity. Although prediction methods like RFNA^26^ and MELD-DNA^27^ can predict a complex from given protein and DNA sequences, they cannot provide insights into binding specificity. In this context, DeepPBS operates on these predicted complexes to yield the binding specificity of the system, which can guide improved modeling of protein-DNA complexes. DeepPBS, despite its generality, exhibits comparable performance to the recently described family-specific method rCLAMPS^28^.

DeepPBS derived RI scores are biologically relevant. They can be aggregated at a protein residue-level, aligning with alanine scanning mutagenesis experimental data. Another advantage of DeepPBS is its speed in predicting binding specificity. Specifically, DeepPBS only requires a single forward call through the model (no need for database search or multiple sequence alignment computation), making it suitable for high-throughput applications like analyzing MD simulation trajectories. In this context, DeepPBS is robust to small dynamical fluctuations and can respond to conformational changes.

The current version of DeepPBS has inherent limitations. It is tailored for double-stranded DNA and is not yet applicable to single-stranded DNA, RNA, or chemically modified bases. However, there is potential for extending the model to accommodate these different scenarios, as well as other polymer-polymer interactions. The DeepPBS architecture can be refined and expanded in terms of applications and engineering enhancements. Collectively, these possibilities hint at an exciting future for molecular interaction studies.

## Methods

### Data sources

The dataset was assembled by integrating protein-DNA structures from the Protein Data Bank (PDB) and their corresponding position weight matrices (PWMs) from JASPAR (2022)^37^ and HOCOMOCO (V11)^31^. These two databases were selected for their accessibility, comprehensive collection, and non-redundancy. The process of merging these three data sources is non-trivial, and detailed description can be found in Supplementary Section 1 and Fig. S1.

### Cross-validation regimen

A 5-fold cross-validation set was constructed with 523 data points as described in Supplementary Section 1. Each datapoint corresponds to a biological assembly containing a protein chain with a corresponding PWM sampled from either JASPAR or HOCOMOCO. The PWM is aligned to DNA in the structure to create a correspondence for loss/metric calculation purposes using an ungapped local alignment process (Supplementary Section 2, *Performance Metrics*). For each fold, cross-validation predictions were made by a model (same for other variations as in Fig. S5a) trained on the remaining four folds (reported in Fig. S5a-c). Full details of training can be found in Supplementary Section 6.

### Benchmark regimen

Datapoints not included in the cross-validation folds were resampled to create a separate benchmark dataset (biological assemblies corresponding to 130 protein chains). This sampling followed the same quality criterion described in Supplementary Section 1, and up to five members per cluster were sampled. Ensemble average predictions of models trained on cross-validation folds are reported for this set in Fig. 2a-d. Combined pre-processing and inference time for one biological assembly is on the order of seconds (e.g., for PDB ID 5X6G, about 15-20 seconds). The DeepPBS ensemble described here was used for all applications on predicted structures.

### Position Weight Matrix (PWM)

For the purposes of this study, a PWM is defined as an NX4 matrix, where N represents the length of the DNA of interest, and the four positions correspond to the four DNA bases: adenine (A), cytosine (C), guanine (G), and thymine (T). Each column in the PWM represents the probabilities of the four bases occurring at that particular position.

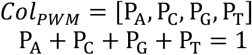

### DNA symmetrization

The DNA representation used is carefully designed with several considerations. Firstly, the DNA sequence in the input complex might not correspond to a high-affinity DNA sequence, particularly in designed structures. Secondly, an all-atom graph representation, similar to the protein, is not convenient because the model ultimately needs to predict a 1D representation (i.e., the PWM) that describes binding specificity. Thirdly, structural data are sparse, and the exact atomic conformation of a bound DNA sequence can make the model overly sensitive and less useful.

Considering these factors, we represent the DNA in a base-symmetrized manner. As shown in Fig. 1c and Fig. S2, this is achieved by designing a symmetrization schema in the base-pair frame, which symmetrizes the seven key atomic interaction positions (4 in major groove, 3 in minor groove). Additionally, four positions are assigned for the sugar and phosphate moieties. For full details of this process, see Supplementary Section 3.

### DeepPBS architecture and training details

Detailed description of the DeepPBS architecture can be found in Supplementary Section 5. Training, cross-validation, and benchmarking details are available in Supplementary Section 6.

### Performance Metrics

Performance metrics used in this work are negative MAE (−*MAE*) and negative RMSE (−R*MSE*), defined as follows:

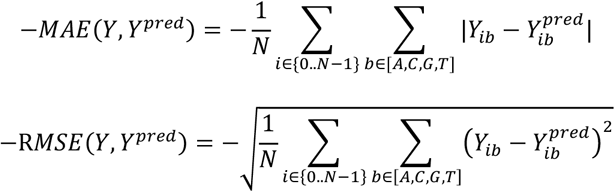

*N* refers to the number of columns in the PWMs being compared. Both metrics follow ‘the higher the better’ principle. They are not independent but have different properties.

### Bipartite edge perturbation and protein heavy atom importance score calculation

Fig. S4 schematically describes the bipartite edge perturbation process for calculating protein heavy atom (say, atom *a*) importance scores. Briefly, the prediction is calculated twice: once (say, *Y*_>_), while considering edges corresponding to the protein heavy atoms, and again (say, *Y*_∼>_) while masking the same edges. This process results in differences in predictions, which can be calculated using the mean absolute difference measure. On their own, these values may not be meaningful, but they can be normalized to the 0-1 range by dividing by the maximum value within a structure. The normalized values, so-called “relative importance” (RI) scores, signify how much the specificity prediction is influenced by interactions made by the corresponding heavy atom. Depending on the downstream use, RI scores can be aggregated at the residue level using either the average, max, or sum aggregations. Mathematically,

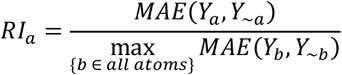

For comparison with alanine scanning mutagenesis experiments (Fig. 4i) at a residue level, the log sum aggregated importance score was calculated. For each atom *a* of a residue *r* in the PDNA interface, let the calculated relative importance be *RI*_>_. Then, this value is calculated as below:

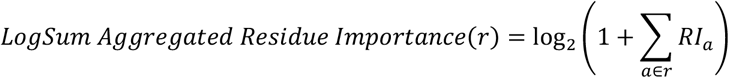

## Data Availability

List of unique identifiers and processed files for cross-validation and benchmark sets, and datasets used for all applicative analysis are available at https://github.com/timkartar/DeepPBS, and preserved as archive at https://doi.org/10.5281/zenodo.8291319.

## Code Availability

Installable source code, pre-trained models, associated guidelines and various custom scripts used for analysis can be found at https://github.com/timkartar/DeepPBS and preserved as archive at https://doi.org/10.5281/zenodo.8291319.

## Supporting information

Supplementary Information: Supplementary Methods, Figures, Table, and References

## Acknowledgements

This work was supported by an Andrew J. Viterbi Fellowship in Computational Biology and Bioinformatics (to R.M.), the National Institutes of Health (grant R35GM130376 to R.R.) and Human Frontier Science Program (grant RGP0021/2018 to R.R.). We acknowledge members of the Rohs lab and Cameron Glasscock (Institute for Protein Design, University of Washington, Seattle, WA) for helpful discussions and comments.

## Author contributions

R.M., J.M.S., and R.R. conceived the project idea with input from T.P.C. R.M., J.M.S., and J.L. designed the model. R.M. and J.M.S. performed data preprocessing. R.M., with input from J.L. and J.M.S., performed model training and benchmarking. R.M., J.L., and T.P.C. developed all application ideas. R.M., J.L., and Y.J. carried out all applications and data analysis. R.M., J.L., Y.J., and R.R. wrote the manuscript. All authors read and commented on the same. R.R. supervised the project.

