## Supplementary Information: Supplementary Methods, Figures, Table, and References for "DeepPBS: Geometric deep learning for interpretable prediction of protein–DNA binding specificity"

1. <u>Datasets</u>

We collected structural data from the Protein Data Bank (PDB)[1] and binding specificity data from JASPAR[2] (version 2022) and HOCOMOCO[3] (v11 core collection). JASPAR catalogues a comprehensive set of experimental binding specificity data for proteins from various species obtained through various types of experiments. HOCOMOCO consists of mainly chromatin immunoprecipitation followed by sequencing (ChIP-seq)[4] experimental data for human and mouse proteins.

Next, we searched for protein-DNA co-crystal data available in the PDB for each position weight matrix (PWM) available to us using corresponding UniProt IDs. We employed DSSR[5] to check for and annotate the existence of one contiguous DNA double helical region in these structures. In our application, we focused on double-stranded DNA only and discarded structures that did not conform to this requirement. Base modifications were replaced by their parent base identity. A total of 1,155 PDB chain IDs were filtered into the dataset. For each structure (biological assembly containing a chain of interest) in the dataset, a corresponding PWM was paired with it. If a PWM existed in both JASPAR[2] and HOCOMOCO[3], one was randomly chosen. PWMs were trimmed to remove uninformative terminal regions with a 0.5 information content (IC) threshold. For each structure, we aligned the corresponding PWM to the DNA helix using an ungapped local alignment (Supplementary Section 2), annotating the region on the DNA helix where predictions should be made and the loss computed during training. For source code and further details of data cleaning and pre-processing, see the Data/Code Availability section.

We clustered the protein chains using CD-HIT[6] with a 40% sequence similarity threshold for clustering, resulting in 189 clusters. This step ensures that our dataset does not overrepresent any particular protein sequence. Next, we sampled up to five members from each cluster, prioritizing biological assemblies where the chain of interest has more contacts with the DNA region where the PWM was aligned. We set the cutoff for alignment length to be at least 5. We split this set of structures into five folds to create a cross-validation set. A schematic representation of this process is shown in Fig. S1a. Experimental and species diversity of the gathered cross-validation dataset are shown in Fig. S1b.

Structures that were not included in the cross-validation dataset were resampled, selecting up to five per cluster following the same criterion. This resulted in 130 datapoints, which we used as a benchmark set. Predictions on this set were only calculated once, after finalizing all models. The family distribution of this set (Fig. 2c) differs from that of the cross-validation set (Fig. S5b).

The PWM of the same protein differed slightly between JASPAR[2] and HOCOMOCO[3] (example shown in Fig. S1c for human estrogen receptor). This observation indicates that there is an inherent limit on what can be possibly learned, signifying noise in collected knowledge. To quantify the performance limit on the dataset based on this phenomenon, we computed the distribution of performance metrics across all unique PWMs appearing in both databases (111 cases).

Alanine scanning mutagenesis involves measuring changes in binding free energy ($\Delta\Delta G$) when performing the same binding experiment for a given protein, with a specific residue mutated to alanine. We used an already gathered dataset[7] of alanine scanning mutagenesis experiments on protein-DNA structures. We filtered the dataset to make it suitable for our context. Specifically, we removed cases involving single-stranded DNA. Mutations to alanine residues with $\Delta\Delta G$ values within 0-3 kcal/mol were retained. We removed cases in which no heavy atom of the mutated residue was within 5 Å of DNA, because our model only assigns importance scores within this range. The final dataset is summarized in Table S1.

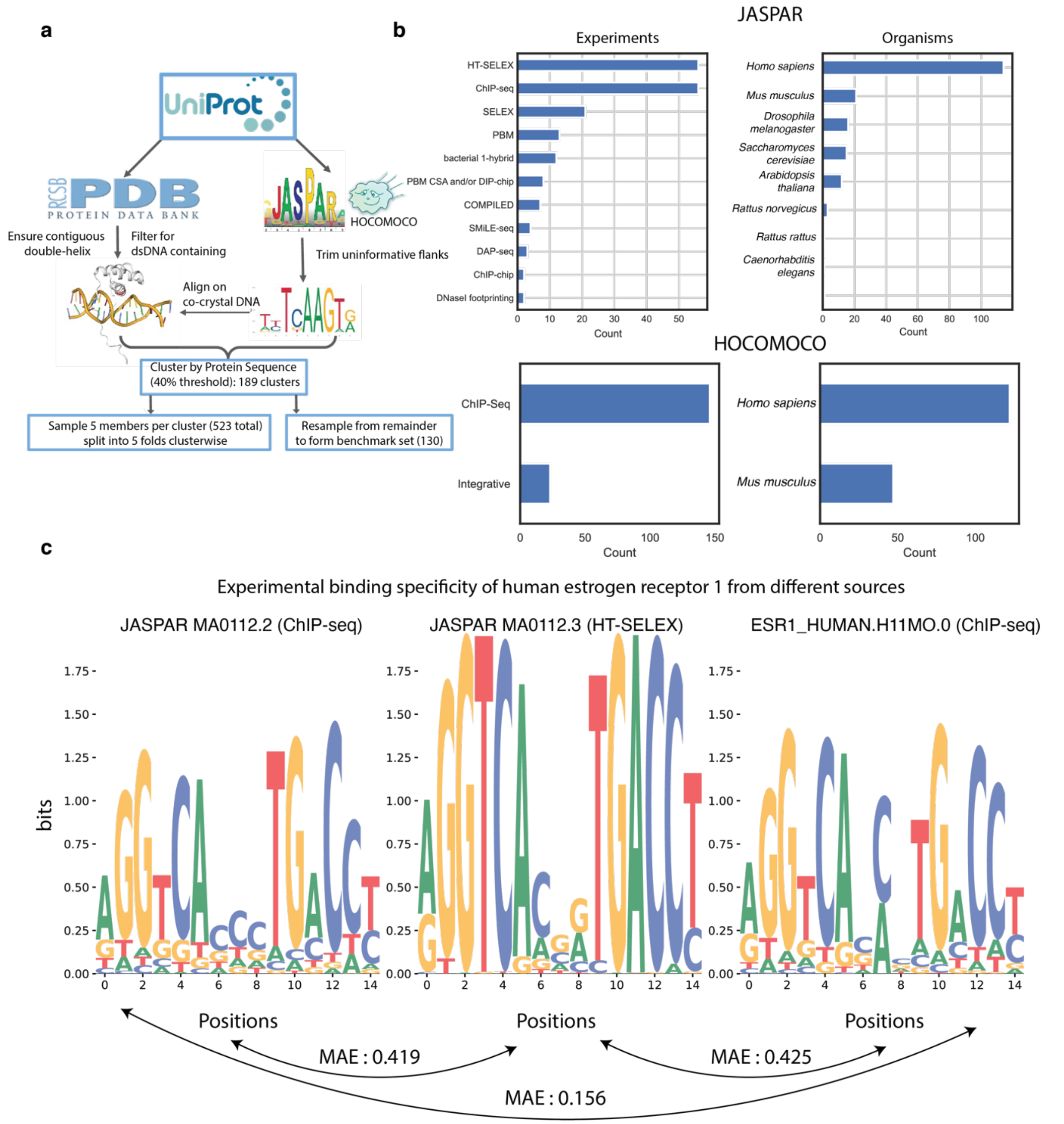

**Fig. S1.**

**Dataset details. (a)** Schematic representation of process for combining data sources. **(b)** Distribution of source experiments and species in constructed cross-validation set. **(c)** Example illustration demonstrating differences in binding specificity data for the same protein (human estrogen receptor 1) from different experiments and databases. Mean absolute error (MAE) over columns among three cases.

### 2. Ungapped Local Alignment

Alignment of experimental PWMs to the corresponding co-crystal DNA is an important step for correctly annotating experimental protein-DNA structural data for model training and evaluation. This alignment needs to be ungapped and should prioritize alignment of higher IC columns from the PWM. Hence, we used an IC-weighted Pearson correlation coefficient (PCC) scoring scheme for the alignment, given by:

$$ICWeightedPCC(Col_{PWM}, Col_{DNA}) \;= PearsonR(Col_{PWM}, Col_{DNA}) \times \frac{1}{2} IC(Col_{PWM})$$

where $PearsonR$ refers to a standard PCC, and IC refers to the information content calculated for a probability simplex with a uniform background, in this context:

$$IC([P_A, P_C, P_G, P_T,]) = \sum_{i \in [A,C,G,T]} \frac{\log(P_i)}{\log(0.25)}$$

<u>Algorithm1 (Pseudocode for ungapped alignment of PWM onto cocrystal DNA using ICWeightedPCC scoring)</u>

```
function ungappedAlign(seq,  pwm):
  // ungapped alignment needs Length X 4 arrays
  max_score ← -9999
  opt_i ←0
  opt_j ←0
  opt_k ←0
  l ← length(seq)
  s ←length(pwm)
  for i from 0 to s-1:
      for k from 0 to s-i:   /// k overlap length
          for j from 0 to l-k:
              score ←0

              for col from 0 to k-1:
                  col_score ←ICWeightedPCC(pwm[i:i+k,:][col,:], seq[j:j+k,:][col,:])
                  score ←score + col_score
              if score > max_score:
                  max_score ←score
                  opt_i ←i //alignment start for PWM
                  opt_j ←j //alignment start for Seq
                  opt_k ←k //alignment length
   return opt_i, opt_j, opt_k, max_score
```

Note, the max_score is length dependent. To compare these values across different PDNA structures (as in Fig. 2b), we have to divide by the overlap length (opt_k).

### 3. Representing DNA

Our framework must consider several important factors for representing DNA. First, our model observes the structure of DNA in the input, but will predict a one-dimensional (1D) representation (a PWM). Thus, from an engineering perspective, it is beneficial to have the same number of features per base pair. Second, the input co-crystal DNA has a sequence; depending on the use case, we may or may not want our model to observe this sequence. Moreover, because experimental structural data are sparse, the co-crystal sequence has a strong potential for overfitting if observed by the model in the input. Therefore, in general, we want to symmetrize each base pair such that all sequence information is lost, but the global shape of the DNA helix is preserved.

With these points in mind, we developed a coarse symmetrized representation of DNA, where each base pair is represented by 11 points: 2 points for the phosphate moiety on each strand, 2 points for the sugar moiety, 4 points for the major groove, and 3 points for the minor groove. Major and minor groove points are placed symmetrically in the base-pair plane, so that they do not possess any particular base identity but roughly correspond to the major and minor groove chemical positions known[8] to be used for base readout. The phosphate moiety is represented by the coordinate of the phosphorus atom. The sugar moiety is represented by the average coordinate of all sugar atoms. The three minor groove points divide the line segment connecting the two C1' atoms into four equal segments. The base-pair plane is determined by the triangle connecting the two C1' atoms and point O (average of atoms N1 and N9). Next, we move perpendicular (to the minor groove line segment) in this plane from either C1' for 3.75 Å and expand the line segment by another 1.54 Å in either direction. The line segment is divided into five equal segments to determine positions of the four major groove points. Additionally, the middle two major groove points are shifted by an additional 1 Å. This geometric construction is based solely on domain knowledge; no learning is employed to estimate any parameter. Fig. S2a shows a schematic representation of this process for an AT base-pair. The only base atoms used for this process are N1 and N9, making it agnostic of base identity.

Fig. S2b shows an example transformation of a DNA structure to a symmetrized helix (sym-helix) using the described process. Fig. S2c shows one CG base pair overlayed with sym-helix points computed for the corresponding base pair. As a result, the DNA structure is represented as $G^d = (V^d, X^d, N^d)$. $V^d$ represents coordinates of the sym-helix points, and $X^d$ represents point-level DNA features, which reflect a one-hot encoded annotation of the 11 positions in the symmetrized base-pair representation. If desired, we can reintroduce the DNA sequence ("DeepPBS with DNASeqInfo" model) by including base-pair-specific chemical group features for each point to $X^d$, as in Chiu et. al. 2023[8]. For each point $v \in X^d$, we also define an interaction vector $N_v^d$. These vectors act as reference directions in the base-pair frame. They are used to compute relative orientation-based features coupled with vectors $N^p$ on the protein graph (refer to Section 4). For the phosphate point, this vector is the average direction of the two double-bonded oxygens; for the sugar point, this vector is the direction of the C4'-C5' bond. For the seven major and minor groove points, these directions are determined by connecting each point to the centroid of the heptagon formed by these points. Fig. S2e shows the arrangement of these vectors on a sym-helix. These directions do not encode any base-specific information and only serve to inform the relative orientation of a sym-helix point in the context of binding. In addition, we include 14 DNA shape features[9,10] denoted as $X^s$, which are base-pair level features (Fig. S2d). These features are: buckle, shear, stretch, stagger, propeller-twist, opening, shift, slide, rise, tilt, roll, helix-twist, major groove width, and minor groove width.

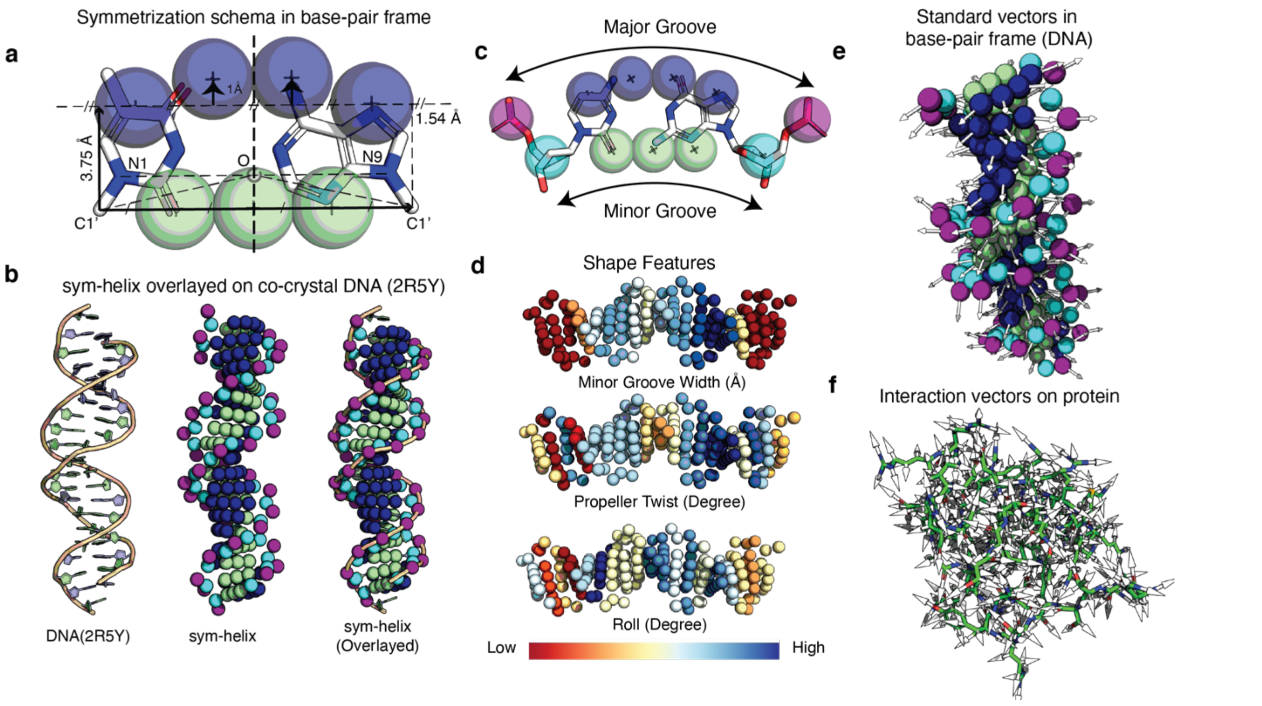

**Fig. S2.**

**Data Representation. (a)** Coarse symmetrization schema at DNA base-pair level. **(b)** Example illustration showing how the computed sym-helix compares with original structure. Each sym-helix point is shown as a sphere (1.5 Å radius) for visibility. **(c)** Symmetrized base-pair representation on one C-G base pair (four major groove points, three minor groove points, and two points each for sugar and phosphate moieties). **(d)** Example computed shape features overlayed on sym-helix as base pair-level features. **(e)** Standard vectors computed on sym-helix, used by the network to correlate orientation information with **(f)** interaction vectors on protein atom-graph, based on average direction of covalent bonds for each heavy atom.

### 4. Representing Protein

In our framework, the protein structure is viewed as a spatial graph $G^p = (V^p, X^p, E^p, N^p)$, where the coordinates of the heavy atoms constitute the vertices $V^p$. For each vertex $v \in V^p$, we define a set of features $X_v^p$ which include the one-hot encoded atom type, solvent-accessible surface area of the atom, charge, radius, circular variance (7.5 Å radius), and Atchley factors[11]. The edges $E^p$ of the protein graph are determined by the covalent bonds; namely, if vertices $u$ and $v$ have a covalent bond between them, then $(u, v) \in E^p$. The edges are unordered. Lastly, to encode directionality of protein side-chain atoms, we encode a unit vector $N_v^p$ for each vertex $v$, computed by averaging the directions of covalent bonds associated with each heavy atom (Fig. S2f).

### 5. DeepPBS Architecture

The architecture of DeepPBS is modular. First, the **ProteinEncoder** module applies spatial graph convolutions on the protein graph to aggregate neighborhood environment information for each protein heavy atom. Initially, a fully connected embedding layer is applied to $X_v^p \; \forall \, v \in G^p$, which expands the dimensionality of $X_v^p$ to 10 dimensions. Four layers of crystal graph convolutions (CGConv)[12] are applied. The first two layers use only covalent bond edges, and the next two layers use distance-based edges with a 4Å radius. The mathematical description of the message-passing scheme for CGConv is as follows:

$$X_v^p \leftarrow X_v^p + \left(\frac{1}{|\mathcal{N}(v)|}\right) * \sum_{u \in \mathcal{N}(v)} \sigma\left(z_{uv} W_f + b_f\right) \odot g(z_{uv} W_s + b_s)$$

where $\mathcal{N}(v)$ denotes the neighbors of $v \in V^p$, and $z_{uv} = [X_v^p, X_u^p, e_{uv}]$ denotes the concatenation of target node features, source/neighboring node features, and edge features (here, the distance between $u$ and $v$). In addition, $\sigma$ denotes the sigmoid function, and $g$ denotes the softplus function. A Rectified Linear Unit[13] (ReLU) activation function is applied after each round of graph convolutions. This marks the end of the ProteinEncoder module.

The sym-helix point features $X_v^d$ are embedded into a 10-dimensional (10D) space using fully connected neural network layers and are used in the next module, the **Bipartite Geometric Network (BiNet)**. In this module, aggregated information on $G^p$ are pulled onto the sym-helix by performing geometry-aware bipartite convolutions. We use a modified version of point pair feature convolutions (PPFConv)[14], which we call a bipartite ResidualPPFConv. The message-passing update scheme associated with it is as follows:

$$X_v^d \leftarrow X_v^d + \gamma_\theta\left(\sum_{u \in \mathcal{N}(v), u \in V^p} h_\theta\left(X_u^p, ||d_{uv}||, \angle(N_v^d, d_{uv}), \angle\left(N_u^p, d_{uv}\right), \angle\left(N_v^d, N_u^p\right)\right)\right)$$

where $d_{uv}$ denotes the line segments connecting a sym-helix point $v$ and a protein heavy atom point $u$. $h_\theta$ is a transformation parametrized by fully connected neural networks. We set $\gamma_\theta$ to be the identity transformation. Four separate ResidualPPFConvs are applied for the major groove, minor groove, phosphate, and sugar points, respectively (from neighbors within 5 Å), followed by ReLU activation. At this stage, we have aggregated all local chemical and geometric interaction contexts onto the sym-helix.

The final module is the **CNN-Predictor** module. We flatten the helix into a 1D base pair-level representation and apply a Multi Layered Perceptron (MLP) to reduce the dimensionality for each base-pair to 32. We concatenate precomputed helix shape features to aggregated base pair-level features. This step allows the network to make connections/correlate patterns between the aggregated information from BiNet and the global shape of the helix. We apply two rounds of 1D convolutions of filter size 3 with a stride of 1. We also apply relevant padding and set output feature size of 8, followed by ReLU activation, making the effective field of view five base-pairs. Next, we apply an MLP for each base-pair to generate logits for predicted base probabilities ($[L_A, L_C, L_G, L_T]$) for corresponding base-pairs. We apply a SoftMax[15] activation to generate the output DNA base probabilities ($[P_A, P_C, P_G, P_T]$). A global temperature parameter ($T_{glob}$) is learned for SoftMax through the training process. Fig. S3 schematically describes the DeepPBS architecture.

$$P_i = \frac{e^{\frac{L_i}{T_{glob}}}}{\sum_{j \in \{A,C,G,T\}} e^{\frac{L_j}{T_{glob}}}} \quad \forall\, i \in \{A, C, G, T\}$$

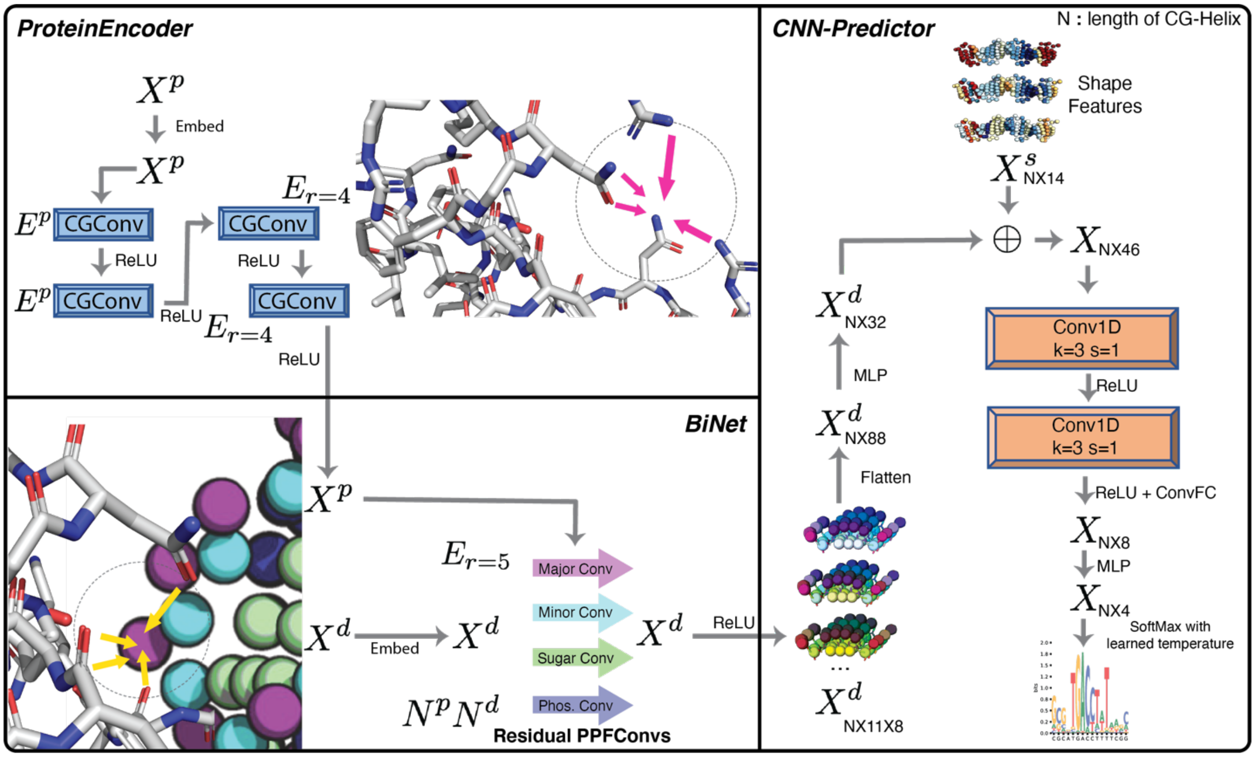

**Fig. S3.**

**DeepPBS architecture.** The DeepPBS architecture can be compartmentalized into three modules: ProteinEncoder, which encodes the protein neighborhood through spatial graph convolutions; BiNet, which consists of a network of bipartite geometric convolutions from the protein graph (same notation as in Supplementary Section 4, $G^p = (V^p, X^p, E^p, N^p)$) to sym-helix points (DNA) (same notation as in Supplementary Section 3, $G^d = (V^d, X^d, N^d)$); and CNN-Predictor, which flattens the aggregated sym-helix features into a 1D representation, adds shape features ($X^s$), and applies 1D convolutional layers followed by fully connected prediction layers. The final logits are converted into probability using a SoftMax activation with a learned temperature parameter. $E_{r=x}$ represents edges determined by vertices/points within a radius of $x$ Å.

### 6. Training, Cross-validation, and Benchmarking Details

Five models were trained for each of the four types: *DeepPBS*, *DeepPBS GrooveReadout*, *DeepPBS ShapeReadout*, and *DeepPBS with DNA SeqInfo*. Each model was trained on four folds of the constructed cross-validation set. Training was conducted for 50 epochs with early stopping on an NVIDIA

RTX A4000 using an Adam[16] optimizer, with a learning rate of 0.001 and weight decay of 0.0001. Hyperparameters were set based on domain knowledge and training curves. For every datapoint, two forward passes were made to account for reverse compliment predictions for both strand directions (with relevant index transformations for input; refer to Data/Code Availability). Outputs were concatenated, and MAE loss was calculated with ground truth (corresponding PWM and its reverse compliment concatenated). Predictions were made on the corresponding fifth/validation fold with each model to gather predictions for all datapoints in the 5-fold dataset. These predictions were used to report metrics in Fig. 4a-h, and Fig. S5a-c.

For benchmarking purposes, ensemble averaged (of the 5 trained cross-validation models) predictions are used Fig. 2a-d. The ensemble is also used for results presented in Fig. 3, 4i, 5, and Fig. S5b,d, S6, S7, S8, and S9.

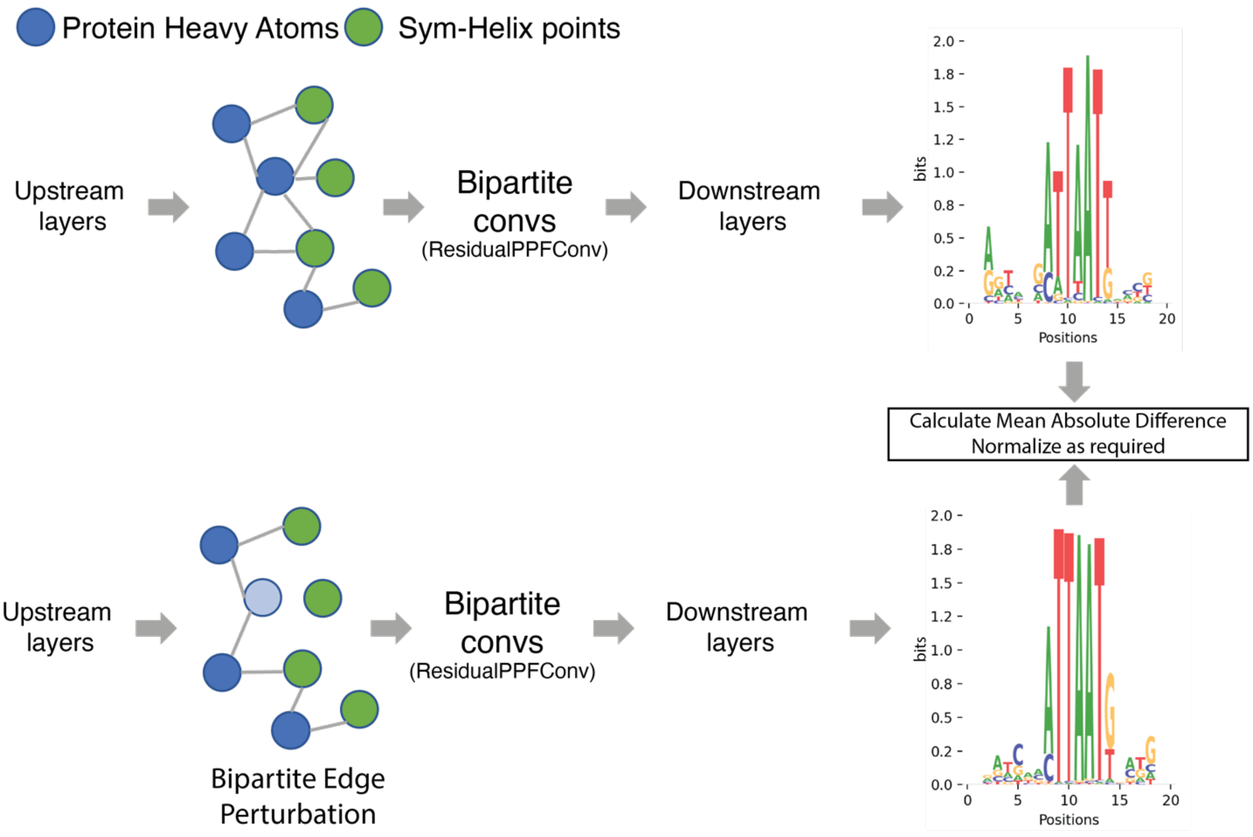

**Fig. S4.**

**Schematic representation of bipartite edge-perturbation process.** Blue circles denote protein heavy atoms. Green circles represent sym-helix points. In one forward pass, the output is calculated with all edges present. In another pass, edges corresponding to one protein heavy atom are excluded from the message-passing scheme, resulting in an alternate output. The difference between the two outputs can be quantified using the mean absolute difference measure and normalized as needed for interpretation purposes.

7. Additional details associated with application of DeepPBS on predicted structures

**Running rCLAMPS:** We ran the rCLAMPS model with default parameters provided by the model's authors according to instructions provided through their GitHub.

**Running RoseTTAfoldNA (RFNA):** We ran the RFNA model with default options and model weights (version: April 13, 2023 v0.2) as presented by the authors through GitHub.

**Trimming unstructured regions from full-length homeodomain (HD) sequences**: For the analysis in Fig. 3e-i, full-length HD sequences were first trimmed to remove unstructured regions, while retaining the main Homeobox domain of interest (rCLAMPS also applies the same process in its pre-processing). This step was achieved by HMMER[17] using the 'homeobox.hmm' file provided by the rCLAMPS repository.

**MM-PBSA vacuum energy calculation**: For each PDB file, we generated a topology file and a run-parameter file using Gromacs 2020.3 to define the force field, amber14sb for protein and parmbsc1 for DNA. These files were used as input for g_mmpbsa to calculate the potential energy in a vacuum. The dielectric constant of solute was set to 8.

**Dataset:** We obtained UniProt protein sequences for three different families, bZIP (homodimers), bHLH (homodimers), and homeodomain (HD) (heterodimers excluded), for cases with a corresponding unique JASPAR entry and no experimental structure for the complex. RFNA predictions that could be successfully processed by DeepPBS pre-processing steps were fed into DeepPBS for specificity prediction (n=49 for bHLH, n=50 for bZIP, n=236 for HD family members).

**Choice of initial guess (IG) DNA:** The IG DNA for the bHLH family was chosen as "GCGCACCACGTGGTGCGC", which has a center E-box motif ("CACGTG") that is known[19] to be a bHLH family target. The IG DNA for the bZIP family was chosen as "GCGCTGATGTCAGCGC" (based on human CREB1 motif MA0018.4). The IG DNA for the HD family was chosen as "GCGTGTAAATGAATTACATGT", based on DNA from PDB ID 1APL.

**Details on metric calculation for Fig. 3d:** We calculated –MAE (best full overlap) for predictions in Fig. 3d against corresponding JASPAR annotations. As a baseline (apart from random predictions drawn from uniform), we calculated –MAE (best full overlap) for the one-hot PWM determined by the IG DNA against the corresponding JASPAR annotation. For the bHLH and HD families, the IG DNA was closer to experimental data than to random baseline (Fig. 3d).

**pLDDT score:** (RFNA-predicted LDDT (Local Distance Difference Test)[20] score). The LDDT score measures the similarity between a predicted and reference structure. When predicting a complex structure, RFNA predicts an LDDT score (pLDDT). These pLDDT scores were shown[21] to be well correlated with the true LDDT of RFNA predictions. Thus, the pLDDT can be taken as a measure of quality of a complex generated by RFNA.

**Comparison of DeepPBS and rCLAMPS**: There are several qualitative advantages to the DeepPBS approach. First, rCLAMPS uses structural mapping for HD-DNA binding to predict specificity for a given HD sequence. This structural mapping leads to a specificity output of exactly 6 base pairs (bp). DeepPBS is functionally not limited to only predicting a 6-bp core and can predict preferences in the flanks (Fig. 3e). Second, rCLAMPS is restricted to predicting monomer preferences (although HD proteins can often bind as dimers; see, e.g., Fig. 5a). In contrast, DeepPBS is able to handle biological assemblies. Third, DeepPBS is not limited to a specific family. For quantitative comparison, we compare the aspects that are achievable by rCLAMPS, namely, 6mer specificity predictions for monomer HD proteins.

**Metric computation for comparison with rCLAMPS:** We ran rCLAMPS on the set of monomer HD proteins (with unavailable complex structure, i.e. not part of RFNA or DeepPBS training). We computed the –MAE (best overlap) values for rCLAMPS predictions against the corresponding JASPAR entry of experimental data, and compared these values to the –MAE values of the best 6-mer overlap for DeepPBS predictions. In this case, for each datapoint, one of round 3, 4 or 5 prediction was chosen. This choice was based on maximizing corresponding protein-DNA contact count (5 Å cutoff) of input RFNA predicted structure.

8. Molecular Dynamics (MD) Simulation of Exd-Scr–DNA system

We conducted MD simulations on the Extradenticle (Exd)-Sex combs reduced (Scr) system, with the dimer bound to its target DNA, using the crystal structure (PDB ID: 2R5Z). AlphaFold2 predictions of the proteins were aligned to the PDB structure to create an initial structure of the simulation. This process aided in filling in the missing linker residues in the biological assembly. The simulation was executed using Gromacs[22] 2020.3 software package. Protein interactions were modeled with the amber14sb[23] force field, and DNA interactions were modeled with the parmbsc1[24] force field. The pdb2gmx program from Gromacs was used to generate topological information for the simulation. The -his flag was used to protonate both Nδ and Nϵ atoms of the His–12 residue for the system with protonated His–12. All complexes were solvated using the explicit TIP3P water model. The negative net charge of the Exd-Hox-DNA complex was neutralized by adding positively charged Na+ counterions, along with negatively charged Cl- counterions, to reach a final NaCl concentration of 150 mM that approximates the physiological concentration. The GROMACS 2020.3 genion program is used to place these counterions throughout the box.

The protein-DNA complex was energy-minimized with steep descent energy minimization for 2,000 steps to relax the structure and remove any steric clashes. Next, we performed three rounds of gradual NVT (constant Number of particles, Volume, and Temperature) equilibration for 10 ps to slowly heat the prepared system to 300K and 1 round of NPT (constant Number of particles, Pressure, and Temperature) to equilibrate the pressure of the system to 1 bar for 700ps. These equilibration rounds were used to adjust the whole system to biological conditions before starting the production simulation. The production simulation for the system was run for 300 ns in the isobaric–isothermal ensemble, where the pressure is maintained at 1 bar and temperature at 300K. The integration time step of 2 fs was used for all calculations. The Verlet cutoff scheme was used for all calculations. Long-range electrostatic interactions were computed using the Particle Mesh Ewald method[25] with a 12Å cutoff. Nonbonded van der Waals interactions were calculated with a 12Å cutoff. The LINCS[26] algorithm was employed to constrain all bonds.

9. Additional details on application of DeepPBS to MD simulation of Exd-Scr–DNA system

The simulation trajectory was divided into 3,000 snapshots (0.1 ns apart), and the DeepPBS ensemble was applied to predict binding specificity for each snapshot. Relative importance (RI) scores were calculated for each heavy atom within 5 Å of DNA, followed by computation of max-aggregated residue RI scores. Fig. S6a shows the initial structure of the simulation, with the locations of some residues of interest marked. In Scr protein, residues Arg5 and His–12 of Scr contribute to minor groove narrowing through electrostatic interactions, which play a crucial role in determining binding specificity[27]. Residues Arg58, Ile57, and Lys61 on the Exd protein interact with the major groove, driving specificity through hydrogen bonding and van der Waals interactions. In the simulation, residues Arg2, Arg3, and Arg5 on Exd contact with the flanking sequences.

Fig. 5b shows the DeepPBS ensemble-predicted PWM, averaged over 3,000 trajectory snapshots. The network demonstrates robustness to dynamic fluctuations throughout the simulation, resulting in an average predicted PWM with a maximum IC nearing 1.5, consistent with the known[28] binding specificity of the system. More interestingly, the network recognizes conformational changes, as reflected by the prediction results and RI scores. Variation of RI of the residues discussed earlier are shown in Fig. 5c and Fig. S6b,d. Movie S1 shows a concurrent view of changes in the network prediction as the simulation progressed, along with corresponding changes in the heavy atom RI score.

Throughout the trajectory, Arg5 and His–12 on Scr consistently interact in the minor groove to drive protein-DNA binding specificity (Fig. S6e). Our model assigns stable RI scores to these residues (Fig. S6d). Arg58B strongly drives specificity by contacting G in the major groove, forming a bidentate hydrogen bond (position 7 in averaged prediction; Fig. 5b). However, after ~100 ns of simulation, the Exd $\alpha$-helix 3 moves closer to the DNA major groove, leading to rotation of Arg58 (Fig. 5c, d) and causing a loss of strong specificity for guanine. Lys61 intermittently contacts the DNA through strong electrostatic interactions, leading to a gain in RI (Fig. 5c, d). On the surface, this kind of importance scoring might appear to replicate some form of distance-based measure. However, our network learns beyond distance relationships. For example, Ile57 remains in close contact with the major groove but only forms van der Waals and other weaker non-electrostatic interactions, resulting in a consistent but lower RI (Fig. 5c, d). This demonstrates the ability of the network to capture more nuanced interactions than simple distance-based measures.

RI scores assigned by our end-to-end deep-learning model offer an efficient alternative to traditional energy calculations, which require meticulous force-field design and energy computations. In the case of residues Arg2, Arg5, and Arg3 at the terminal loop region of the Exd protein, temporal changes in RI scores (Fig. S6b) strongly correspond to conformational changes of these residues over the simulation trajectory, as highlighted in Fig. S6c. Arg2 forms a bidentate hydrogen bond with G (~40 ns to 100 ns), which appears in DeepPBS predictions as highly specific for C (Fig. S6b). Arg5 interacts with an adjacent minor groove for most of the trajectory; however, it deviates away from the minor groove after ~210 ns, and a corresponding reduction in RI is observed. This demonstrates the ability of our deep-learning model to capture the dynamic behavior of residues and their interactions with the DNA.

In summary, DeepPBS has demonstrated its robustness and adaptability in response to both small dynamical fluctuations and conformational changes. Although the model was trained on snapshot structures and experimental PWMs, its predictions and RI scores are well-regularized and versatile, making it suitable for automated analysis of MD trajectories and designed protein-DNA complexes. These factors make DeepPBS a valuable tool for researchers working in the field of protein-DNA interactions, enabling deeper understanding and insights into the behavior of these complex molecular systems.

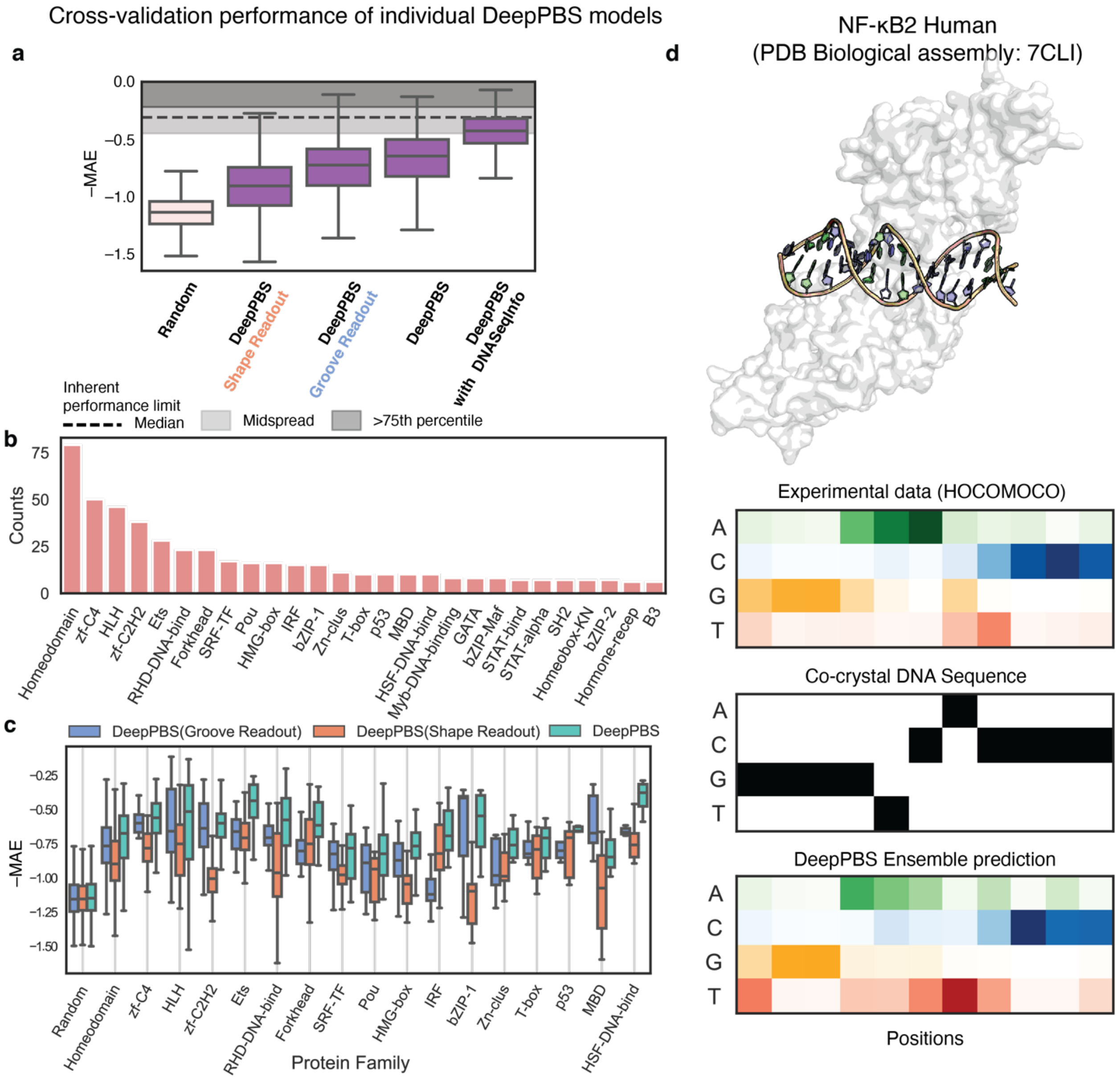

**Fig. S5.**

**Cross-validation performance of DeepPBS for predicting binding specificity across protein families on experimentally determined structures. (a)** Cross-validation performance of individual trained models for the DeepPBS model and its variations, with outliers removed. **(b)** Abundance of various protein families (PFAM annotations) in cross-validation dataset (counts > 5). **(c)** Cross-validation performance of DeepPBS, along with "Groove Readout" and "Shape Readout" variations, across protein families (counts > 8), with outliers removed. **(d)** Example DeepPBS ensemble prediction on NF-κB biological assembly (sampled in benchmark set) containing non-optimal DNA sequence. DeepPBS ensemble prediction on NF-κB biological assembly for human (NFKB2, UniProt ID Q00653) is shown in the benchmark dataset. Although the co-crystal DNA sequence is not of the highest affinity (judging by experimental data from HOCOMOCO), our prediction can circumvent this issue and predict binding specificity levels that are much closer to those observed experimentally. For box plots in **(a)** and **(c)**, lower limit represents lower quartile, middle line represents median, and upper limit represents upper quartile. Whiskers do not include outliers.

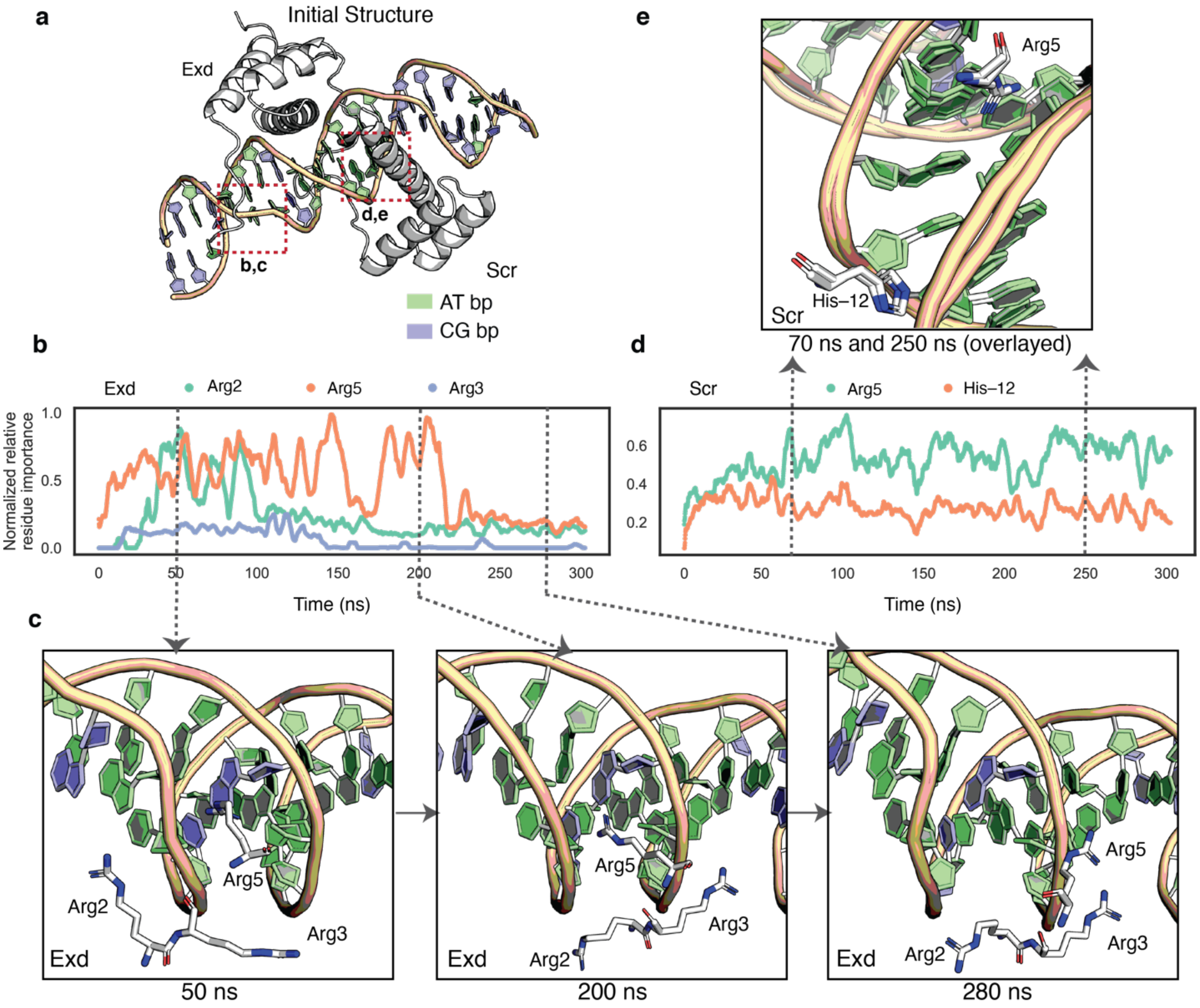

**Fig. S6.**

**Application of DeepPBS to MD simulation of AlphaFold2- and PDB (2R5Z)-based modeled complex of Exd-Scr system. (a)** Initial structure of simulation. Locations of residues of interest are marked. **(b)** Residue-level RI score over time (averaged per 5 ns window) for residues involved in Exd-DNA flank ineraction. **(c)** Snapshot of interactions by Exd Arg2, Arg3, and Arg5 at 50 ns, 200 ns, and 280 ns, respectively. **(d)** Residue-level RI score over time (averaged 5 ns window) for residues involved in Scr-DNA minor groove interaction. **(e)** Snapshot of interactions by Scr Arg5 and His–12 at 70 ns and 250 ns, respectively.

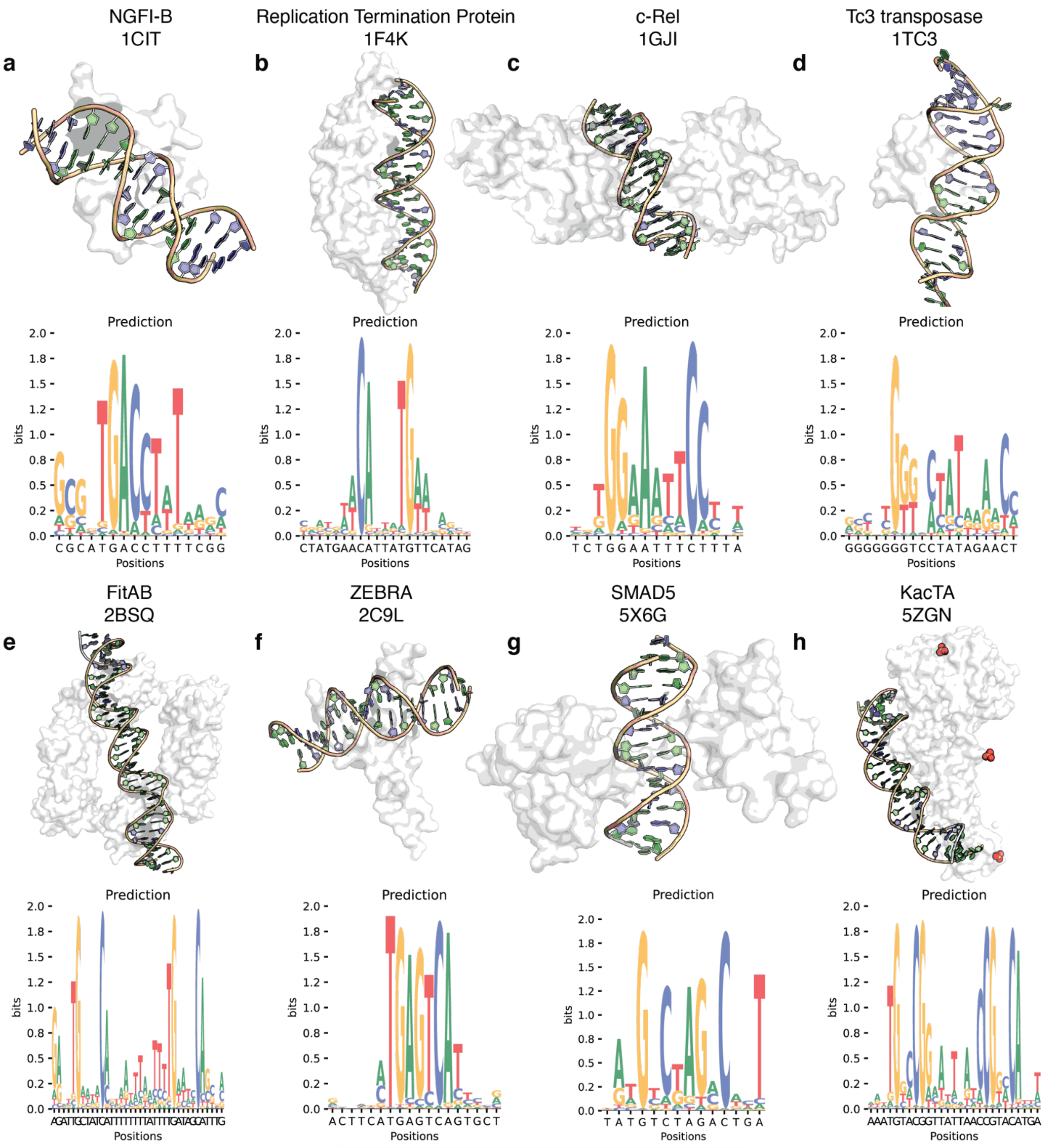

**Fig. S7.**

**Example DeepPBS ensemble predictions on structures of specific binders.** Specificity data were unavailable on JASPAR/HOCOMOCO. **(a)** Monomeric orphan nuclear receptor NGFI-B, **(b)** replication termination protein in bacteria, **(c)** proto-oncogene product c-Rel, **(d)** Tc3 transposase bound to transposon DNA, **(e)** Fitab protein from *Neisseria Gonorrhoeae*, **(f)** Epstein-Barr virus ZEBRA protein, **(g)** Smad5-MH1 protein/ palindromic SBE DNA complex, and **(h)** DUF1778 domain-containing kacTA protein.

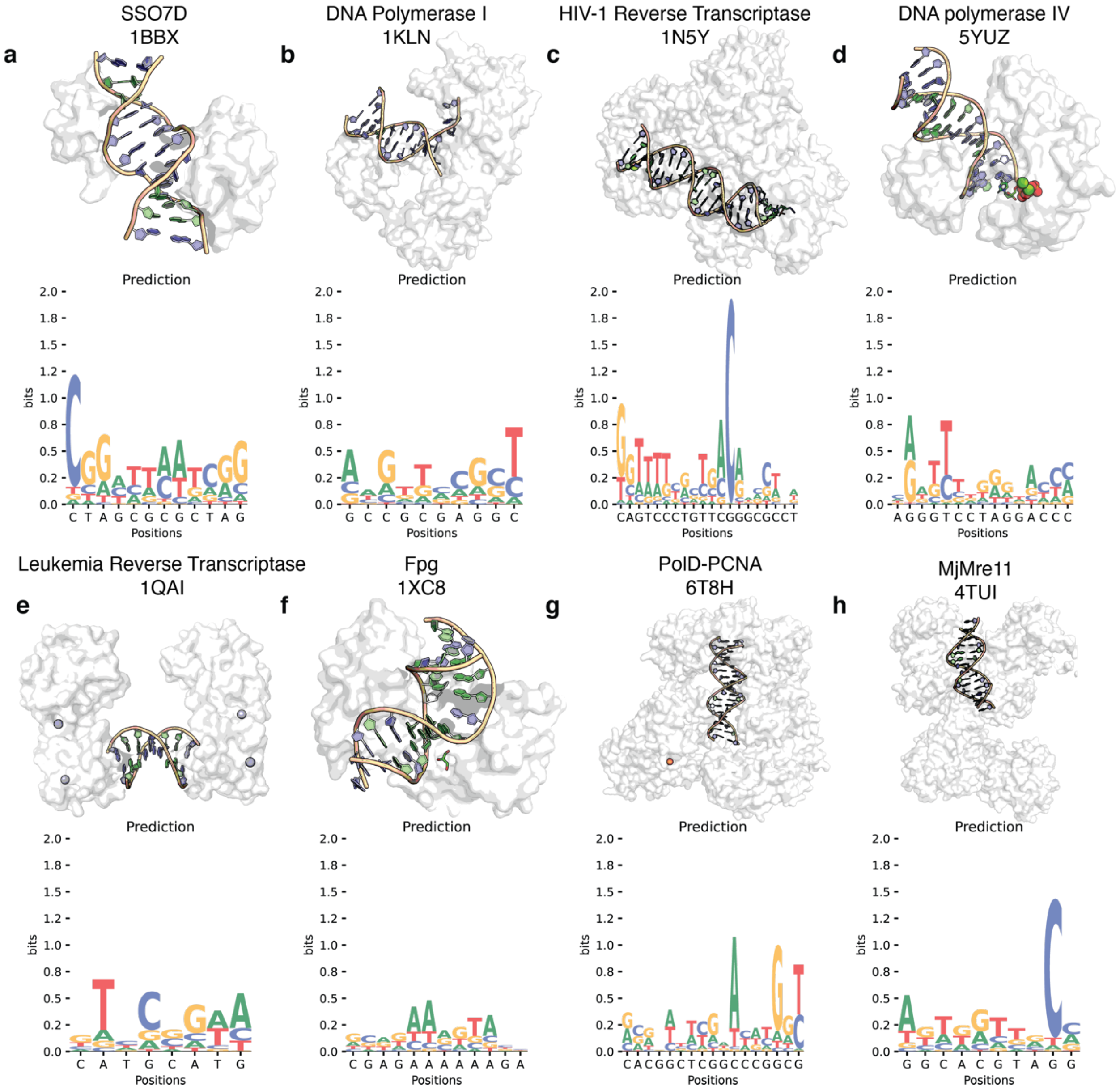

**Fig. S8.**

**Example DeepPBS ensemble predictions on structures of non-specific binders to DNA double helix. (a)** Sso7D-DNA complex, **(b)** DNA polymerase I Klenow fragment, **(c)** HIV-1 reverse transcriptase with pre-translocation and post-translocation AZTMP-terminated DNA, **(d)** DNA polymerase IV, **(e)** Moloney murine leukemia virus reverse transcriptase, **(f)** DNA repair enzyme formamidopyrimidine-DNA glycosylase (Fpg), **(g)** DNA polymerases in complex with proliferative cell nuclear antigen (PCNA), and **(h)** DNA damage sensing enzyme *Methanococcus jannaschii* Mre11 (MjMre11).

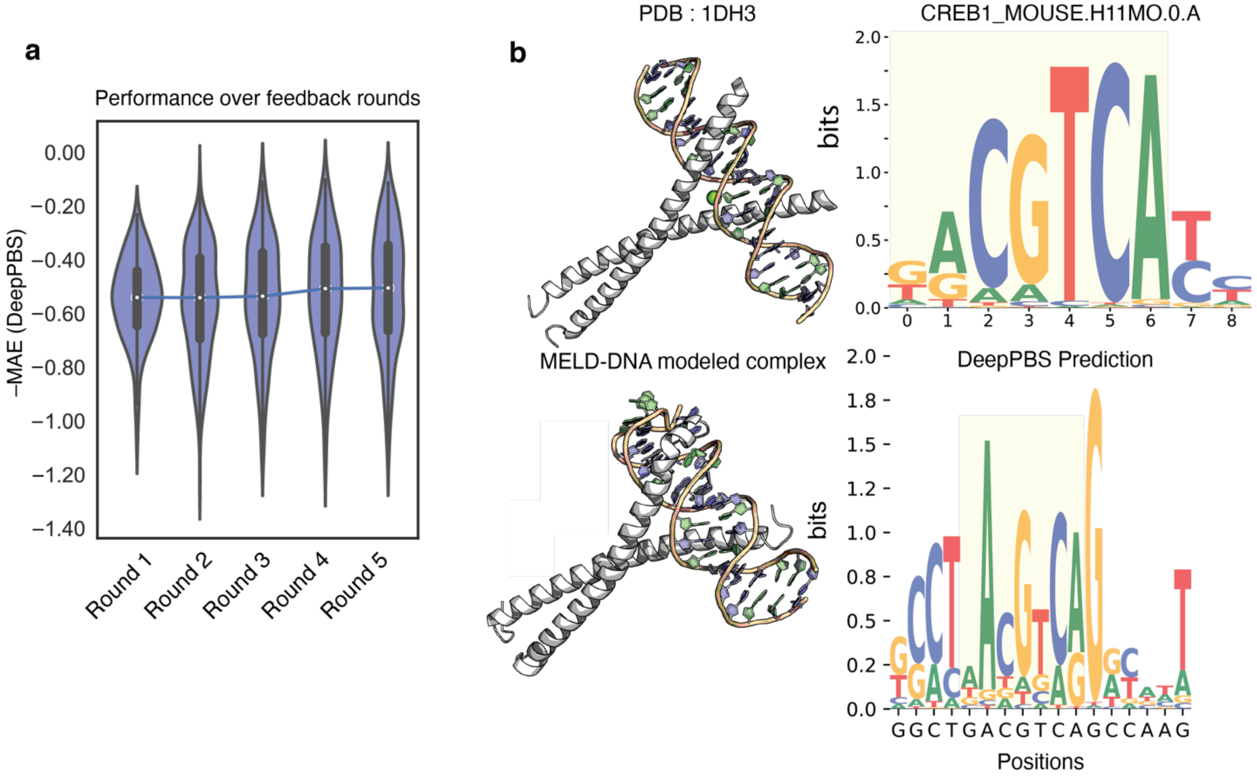

**Fig. S9.**

**Application of DeepPBS on modeled structures. (a)** Performance of DeepPBS (best 6mer overlap) improves as the feedback loop progresses (corresponding to Fig. 3f-g) in conjunction with improvement in RFNA complex design. **(b)** Example application of DeepPBS on mouse CREB1 dimer bound to DNA modeled by MELD-DNA (as provided by authors).

| DeepPBS | ΔΔ$G$ (kcal/mol) | PDB ID | Residue |
|---|---|---|---|
| 2.9589 | 2.62 | 1b3t | Y518 |
| 0.1869 | 0.17 | 1fos | K148 |
| 0.146 | 0.17 | 1fos | K153 |
| 0.1333 | 0.92 | 1fos | R144 |
| 3.3883 | 1.04 | 1fos | R155 |
| 0.1765 | 0.34 | 1fos | R158 |
| 0.1001 | 0.04 | 1fos | R272 |
| 3.0263 | 0.84 | 1hcq | E25 |
| 0.8854 | 0.92 | 1hcq | H18 |
| 4.1678 | 1.21 | 1hcq | K28 |
| 3.1504 | 1.24 | 1hcq | K32 |
| 1.0881 | 1.22 | 1hcq | Y19 |
| 1.1315 | 0.7 | 1j5n | K22 |
| 0.6752 | 0.5 | 1j5n | K53 |
| 1.2293 | 0.4 | 1j5n | K60 |
| 1.0536 | 0.4 | 1j5n | K78 |
| 0.8865 | 0.2 | 1j5n | K85 |
| 3.6399 | 0.5 | 1j5n | M29 |
| 1.1345 | 0.7 | 1j5n | N33 |
| 1.0316 | 0.8 | 1j5n | R36 |
| 0.5118 | 0.7 | 1j5n | R40 |
| 3.4768 | 0.9 | 1j5n | Y28 |
| 0.7347 | 0.2 | 1j5n | Y81 |
| 1.636 | 0.2 | 1mse | S187 |
| 1.5845 | 0.7 | 1tn9 | K21 |
| 3.6307 | 1.36 | 1tn9 | K28 |
| 0.7296 | 1.33 | 1tn9 | K54 |
| 3.5022 | 0.43 | 1tn9 | R20 |
| 1.6292 | 1.22 | 1tn9 | R24 |
| 2.7009 | 1.17 | 1tn9 | R55 |
| 0.1072 | 0.75 | 1tn9 | R5 |
| 0.8861 | 0.04 | 1tn9 | T15 |
| 0.8219 | 0.44 | 1tn9 | W42 |
| 2.934 | 1.51 | 1tn9 | Y40 |
| 0.7094 | 0.2725 | 2mxf | K102 |
| 0.8308 | 0.6225 | 2mxf | K105 |
| 0.8877 | 0.7518 | 2mxf | K108 |
| 0.2503 | 0.283 | 2mxf | K81 |
| 1.8955 | 0.2967 | 2mxf | K97 |
| 4.307 | 0.7725 | 2mxf | N100 |
| 4.2766 | 1.3382 | 2mxf | R80 |
| 4.0195 | 2.0467 | 3ufd | R46 |
| 0.6758 | 0.971 | 3ufd | S52 |
| 2.5213 | 1.4177 | 3ufd | Y37 |
| 3.1102 | 2.2518 | 4bnc | R391 |

**Table S1.**
**Data presented in Fig. 4i along with corresponding PDB IDs and target residues.**

Click here to access
supplementary movie s1

**Movie S1.**

**Concurrent view of changes in network prediction as simulation progressed, along with corresponding changes in heavy atom importance score.**
